# TLR2 Supports γδ T cell IL-17A Response to ocular surface commensals by Metabolic Reprogramming

**DOI:** 10.1101/2024.04.01.587519

**Authors:** Wenjie Zhu, Xiaoyan Xu, Vijayaraj Nagarajan, Jing Guo, Akriti Gupta, Zixuan Peng, Amy Zhang, Jie Liu, Mary J. Mattapallil, Yingyos Jittayasothorn, Reiko Horai, Yasmine Belkaid, Michael Constantinides, Anthony J. St. Leger, Rachel R. Caspi

## Abstract

The ocular surface is a mucosal barrier tissue colonized by commensal microbes, which tune local immunity by eliciting IL-17 from conjunctival γδ T cells to prevent pathogenic infection. The commensal *Corynebacterium mastitidis* (*C. mast*) elicits protective IL-17 responses from conjunctival Vγ4 T cells through a combination of γδ TCR ligation and IL-1 signaling. Here, we identify Vγ6 T cells as a major *C. mast*-responsive subset in the conjunctiva and uncover its unique activation requirements. We demonstrate that Vγ6 cells require not only extrinsic (via dendritic cells) but also intrinsic TLR2 stimulation for optimal IL-17A response. Mechanistically, intrinsic TLR2 signaling was associated with epigenetic changes and enhanced expression of genes responsible for metabolic shift to fatty acid oxidation to support *Il17a* transcription. We identify one key transcription factor, IκBζ, which is upregulated by TLR2 stimulation and is essential for this program. Our study highlights the importance of intrinsic TLR2 signaling in driving metabolic reprogramming and production of IL-17A in microbiome-specific mucosal γδ T cells.

**Summary:** Ocular commensal *Corynebacterium mastitidis* (*C. mast*) induces IL-17 responses from γδ T cells by activating TLR2 signaling. γδ T cell-intrinsic TLR2 stimulation can increase *Il17a* transcription and promote fatty acid oxidation, favoring γδ T cell IL-17A responses.

**Highlights:**

(1) Vγ6 T cells are a major *C. mast*-responsive subset in the conjunctiva
(2) TLR2-deficient mice exhibit reduced γδ T cell responses to ocular commensal bacteria.
(3) γδ T cell-intrinsic TLR2 deficiency causes defects of fatty acid oxidation and IL-17A production in a γδ subset-specific manner.
(4) The transcription factor, IκBζ is upregulated by TLR2 stimulation and supports γδ IL-17A production through fatty acid oxidation.

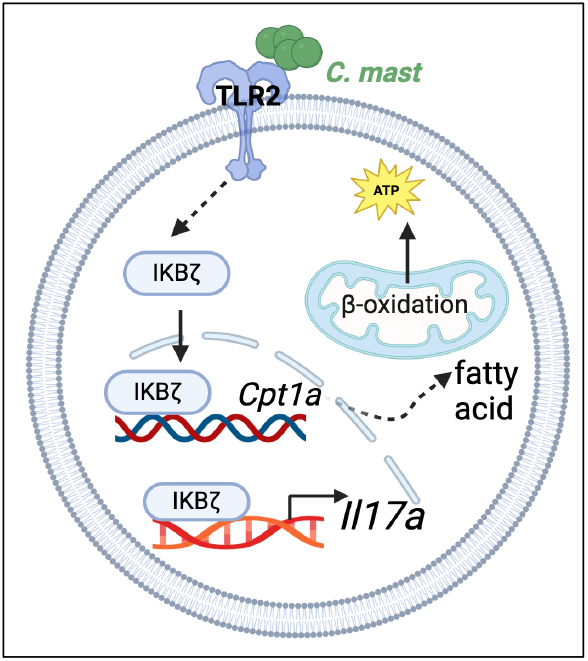

## INTRODUCTION

An immunological balance between microbes and mucosal barrier tissues is required to ensure tissue health and homeostasis. Specifically, immune cells must mount a response that limits the outgrowth of microbes from becoming pathobionts without causing immunopathology. The cytokine interleukin (IL)-17 plays a critical role in this process. IL-17 is required at virtually every mucosal site in the body to limit outgrowth of bacteria and fungi in the steady-state^1^. Humans and mice deficient in production of IL-17A and F experience severe and life-threatening infections^2^.

Regulation of IL-17 production in CD4+ αβ T cells relies on a combination of T cell receptor stimulation and signals from proinflammatory cytokines like IL-23, IL-1β, and IL-6^3,4^. Regulation of IL-17 production in unconventional lymphoid cells, such as invariant natural killer T cells (iNKT), innate lymphoid cells (ILC), and γδ T cells, which often have critical biological functions at mucosal barrier sites, is less well-understood. Although IL-1β signaling is required by iNKT and γδ T cells for IL-17 production^5^, TCR ligation is essential for an optimal response^6^. We as well as others have demonstrated the critical role of IL-17 produced by unconventional T cells in autoimmunity and mucosal host defense, highlighting the importance of mechanisms that govern the effector functions of these cells^7-9^.

The conjunctiva of the eye is a specialized mucosal tissue responsible for protecting the ocular surface against allergic and infectious insults^10^. Mice deficient in IL-17A and IL-17F are highly susceptible to severe bacterial and fungal infections, including corneal *Candida albicans* infection and the spontaneous outgrowth of *Staphylococcus spp*.^7,11^. We previously demonstrated that conjunctival γδ T cells are key contributors to the protective IL-17 elicited at the ocular surface by the Gram-positive commensal bacterium, *Corynebacterium mastitidis (C. mast)* ^12^. In Vγ4 γδ T cells, IL-17A production relied on TCR ligation and IL-1β signaling. Notably, *C. mast* also stimulated IL-17A production in Vγ4-negative γδ T cells, but their identity and the signals they required to induce IL-17A were unclear. Tight control over the production of conjunctival IL-17A is required to both prevent the outgrowth of pathobionts, and to limit bystander damage to the ocular surface, such as occurs in dry eye and other ocular surface diseases^13-15^. Therefore, understanding how an eye-colonizing microbe stimulates IL-17 in the Vγ4-negative γδ T cell subset will be essential for deciphering how mucosal immune responses are maintained at the ocular surface.

IL-17 responses in γδ T cells can be stimulated indirectly by microbial components which are sensed by toll-like receptors (TLRs) on myeloid cells that produce IL-1β and IL-23, which in turn stimulate IL-17A-producing γδ Τ cells (γδΤ17)^16^. However, γδ Τ cells themselves express various TLRs, including TLR2, TLR4, TLR6 and TLR9, which can affect their effector functions. For example, direct TLR2 engagement *in vivo* by the synthetic LPS analog Pam3CSK4 enhances γδ T cell proliferation and IL-17 production^17-19^.

In addition to immune activation pathways, the metabolic regulation of γδΤ17 cells has recently become an area of intense interest. The development and the pro-tumor function of γδΤ17 cells were shown to be under metabolic control^20,21^. However, the molecular mechanisms and the role of TLR2 in metabolic programming of these cells are not known. Our study offers to bridge these gaps by elucidating how TLR2 signaling influences IL-17 production through metabolic reprogramming of *C. mast*-specific γδ T cells.

Utilizing TLR2-deficient (TLR2−/−) mice colonized with *C. mast*, we demonstrate that TLR2 expression is essential in both dendritic cells (DCs) and in γδ T cells themselves for optimal IL-17A production. Notably, intrinsic TLR2 expression occurred mainly in Vγ6 cells, which we identify here as the major conjunctival γδ Τ cells and key responders to *C. mast*. Furthermore, our findings reveal that intrinsic TLR2 signaling in Vγ6 γδ T cells is linked to activation of the transcription factor IκBζ and causes a metabolic shift to fatty acid oxidation to support IL-17 production. Our findings underscore the significance of TLR2 in the metabolic and immune regulatory pathways of mucosal γδ T cells and highlight TLR2 as a potential target for modulating immune responses at mucosal barriers.

## RESULTS

### TLR2 is required for IL-17A production in response to *C. mast* at the ocular surface

Ocular colonization with the microbe, *C. mast*, results in the expansion of γδΤ17 cells in the eye draining lymph nodes (DLNs)^12^. These cells accumulate in the conjunctiva where their functionality leads to a steady-state, non-pathogenic recruitment of neutrophils to the tissue. Even though TCR stimulation and IL-1β were shown to induce IL-17, the requirement of innate receptors in the production of IL-17 in these cells was still unresolved. Therefore, we focused our attention on the requirement of TLR2 signaling in mucosal γδ T cell IL-17 responses. We ocularly colonized WT or TLR2^−/−^ mice with *C. mast*, as previously reported, and assessed the production of IL-17A in DLNs. We found the percentage of IL-17A positive cells and the mean fluorescence index (MFI) of IL-17A in DLNs were reduced in TLR2^−/−^ mice compared to WT mice (Fig. 1A - B). This was due, in part, to a reduction in proliferation as measured by the expression of Ki67 within DLN γδ T cells (Fig. 1C). There was a concurrent reduction of IL-17 in the γδ T cell compartment of the conjunctiva (Fig. 1D). The effect of the reduction in IL-17 was also observed in the recruitment of neutrophils to the conjunctiva (Fig. 1E).

**Figure 1.**
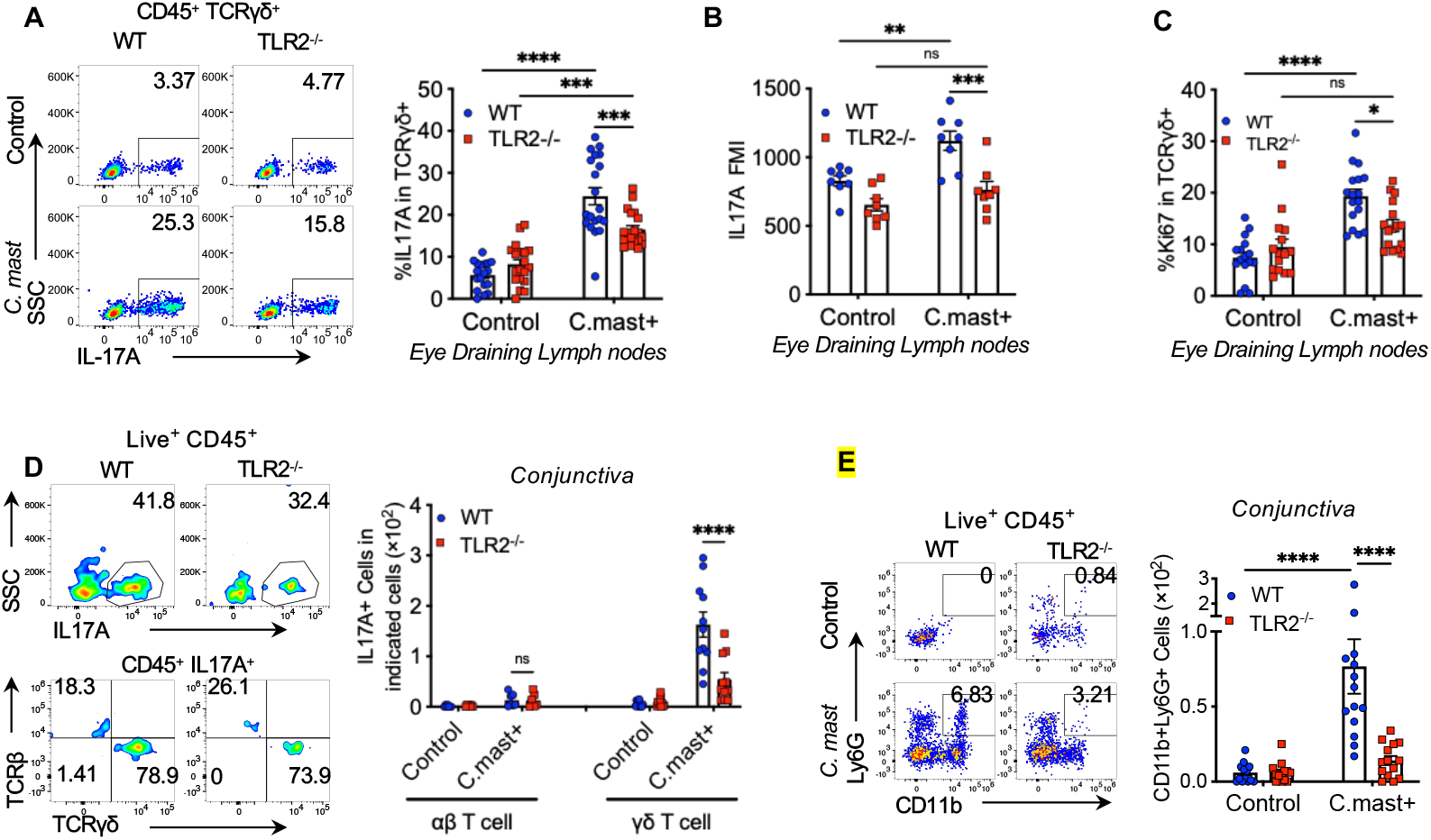
TLR2 is required for IL-17A production in response to *C. mast* at the ocular surface. *C. mast* (10^8^ CFU) or PBS was instilled onto each eye every 3 days for a total of 3 treatments. **(A)** Representative FACS plots and bar graphs showing the percentage of IL17A^**+**^ in γδ T cells from eye-draining lymph nodes. N= 20. Combined data from 6 experiments. **(B)**Bar graphs depict geometric mean fluorescence intensity (MFI) of IL17A in eye-draining lymph node γδ T cells. N= 8. Combined data from 2 experiments **(C)**The percentage of Ki67+ γδ T cells in eye draining lymph node. N=15-17. Combined data from 5-6 experiments. **(D)**Representative FACS plots showing the percentage of IL-17A-producing cells (top panels), IL17A+TCRαβ+ cells, and IL-17A+TCRγδ+ cells in the conjunctiva (bottom panels). Bar graphs show the number of conjunctival IL17A+TCRαβ+ cells and IL-17A+TCRγδ+ cells. IL17A+TCRαβ+ cells: data combined from 2 experiments. N= 7. IL-17A+TCRγδ+ cells: data combined from 5 experiments. N=14 per group. **(E)**Representative FACS plots and bar graphs showing the cell number of conjunctival neutrophils in WT and TLR2^−/−^ mice with/without *C. mast* inoculation. N=14. Combined data from 5 experiments. (A-E) Bars represent mean ± SEM with *P<0.05, **P<0.01, ***P <0.001, ****P<0.0001. Statistical significance was determined by two-way ANOVA (A-E).

### TLR2 on γδ T cells is required for their IL-17A response to *C. mast in vitro*

TLR2 is expressed on γδ Τ cells and DCs at inflammatory sites ^22,23^ and is required for microbe-specific IL-17 production, we sought to identify the immune cells that express TLR2 in the context of ocular *C. mast* colonization. Within the conjunctiva, both DCs and γδ T cells express detectable levels of TLR2 after *C. mast* inoculation, suggesting a role for this receptor in *C. mast*-specific IL-17 response (Fig. 2A). In contrast, conjunctival αβ T cells did not express detectable levels of TLR2 regardless of the presence of *C. mast*.

**Figure 2.**
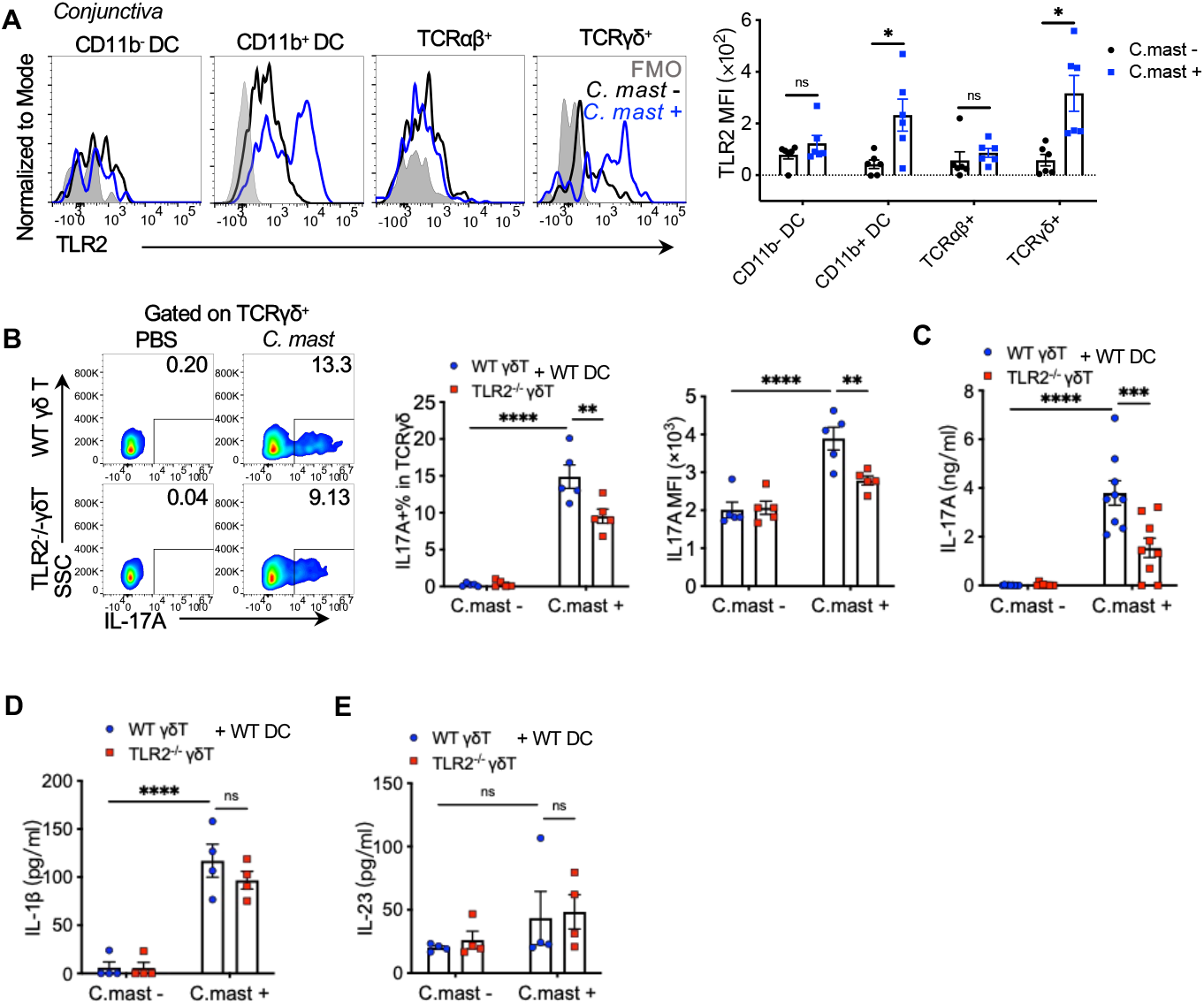
TLR2 expression in γδ T cells is needed for full IL-17A Responses to *C. mast*. WT or TLR2^−/−^ γδ T cells were sorted by flow cytometry from naive mice and co-cocultured with WT CD11c^+^ dendritic cells for 72 hours (γδ T cells: 2×10^4^; CD11c+ cells: 1×10^5^) with/ without heat-killed *C. mast*. Supernatants and cells were collected for ELISA or flow cytometry analysis. **(A)** Representative FACS plots and bar graphs depicting TLR2 expression in CD11b^-^ DC, CD11b^+^ DC, TCRαβ^+^ and TCRγδ^+^ cells in the conjunctiva of C57BL/6J WT mice with or without *C. mast* association. N = 5 per group. Combined data from 2 experiments. **(B)**Representative FACS plots and bar plots showing the percentage and MFI of IL-17A on γδ T cells from WT or TLR2^−/−^ γδ T cell co-culture system. N= 5. Combined data from 3 experiments. **(C-E)** Bar graphs presenting the levels of IL-17A **(C)**, IL-1β **(D)**, and IL-23 **(E)** in the supernatants from WT or TLR2^−/−^ γδ T cell co-culture system. Data was combined from 4 experiments (N= 9) for (C); and 2 experiments (N= 4) for (D-E). (A-E) Bars represent mean ± SEM with *P<0.05, **P<0.01, ***P <0.001, ****P<0.0001. Statistical significance was determined by Welch’s t-test (A) or two-way ANOVA (B-E).

These data prompted the question of whether TLR2 expression is necessary on DCs, γδ T cells, or both for *C. mast*-specific IL-17 response. To address this, we examined co-cultures of WT and TLR2-deficient DCs and γδ T cells in all reciprocal combinations, in the presence or absence of heat-killed *C. mast*. Stimulation with *C. mast* expectedly enhanced IL-17A production in cultures containing WT DCs and WT γδ T cells. In contrast, TLR2^−/−^ DCs supported a significantly reduced γδ IL-17 response: fewer IL-17A+ γδ T cells, lower MFI of IL-17-positive cells (Fig. S1A), and less secreted IL-17A as measured by ELISA (Fig. S1B). Because DCs activated by TLR2 signals secrete IL-1β and IL-23, needed for γδ T cells to express IL-17A, we measured the levels of these cytokines in the co-culture supernatants. TLR2^−/−^ DCs produced less IL-1β (but not IL-23) in response to *C. mast* stimulation (Fig. S1C-D), suggesting that diminished IL-1β production by TLR2^−/−^ DCs may have accounted for the reduced IL-17A production by γδ Τ cells. This was confirmed by the ability of exogenous IL-1β to restore IL-17A production to levels comparable to those observed with WT DCs (Fig. S1E).

Next, we examined the effect of TLR2 deficiency in γδ T cells in their response to *C. mast*. In the presence of WT DCs, TLR2^−/−^ γδ Τ cells still produced less IL-17A than WT, by flow cytometry and ELISA (Fig. 2B-C), despite undiminished IL-1β and IL-23 (Fig. 2D-E). This indicated that optimal IL-17A production in response to *C. mast* in these cultures required TLR2 expression on both DCs and γδ T cells. While the need for TLR2 on DCs can be explained by their production of IL-1β, endogenous TLR2 expression on γδ Τ cells is independently needed to support their IL-17 response.

### The absence of TLR2 on γδ T cells decreases their IL-17A responses *in vivo*

To study the γδ cell-intrinsic role of TLR2 *in vivo*, we performed adoptive transfer of TLR2-sufficient (WT) or deficient (KO) γδ T cells into TCRδ^−/−^ mice, which lack γδ T cells entirely. Donor γδ Τ cells were purified by sorting from (*C. mast*–naïve) CD45.1 WT or CD45.2 TLR2^−/−^ mice and were co-transferred at a 1:1 ratio to sublethally irradiated (450 R) TCRδ^−/−^ recipient mice. One week after cell transfer, the recipients were ocularly associated with *C. mast* or PBS as a control, and after another week, we assessed their IL-17A responses (Fig. 3A). In the *C. mast*-unassociated group, both WT and TLR2^−/−^ γδ T cells populated the recipient mice with similar efficiency, indicating that TLR2 is not required for γδ Τ cell “fitness” over the observation time period (Fig. 3B). However, in mice associated with *C. mast*, TLR2^−/−^ γδ Τ cells displayed reduced responses compared to the co-transferred WT γδ T cells (Fig. 3B). We observed a lower percentage of IL-17A positive cells, diminished ex-vivo IL-17A production to PMA per cell (MFI), and a lower proportion of proliferating (Ki67^+^) cells (Fig. 3C-D). These results confirm the cell-intrinsic role of TLR2 on γδ T cells for activation and IL-17A production in response to *C. mast in vivo*.

**Figure 3.**
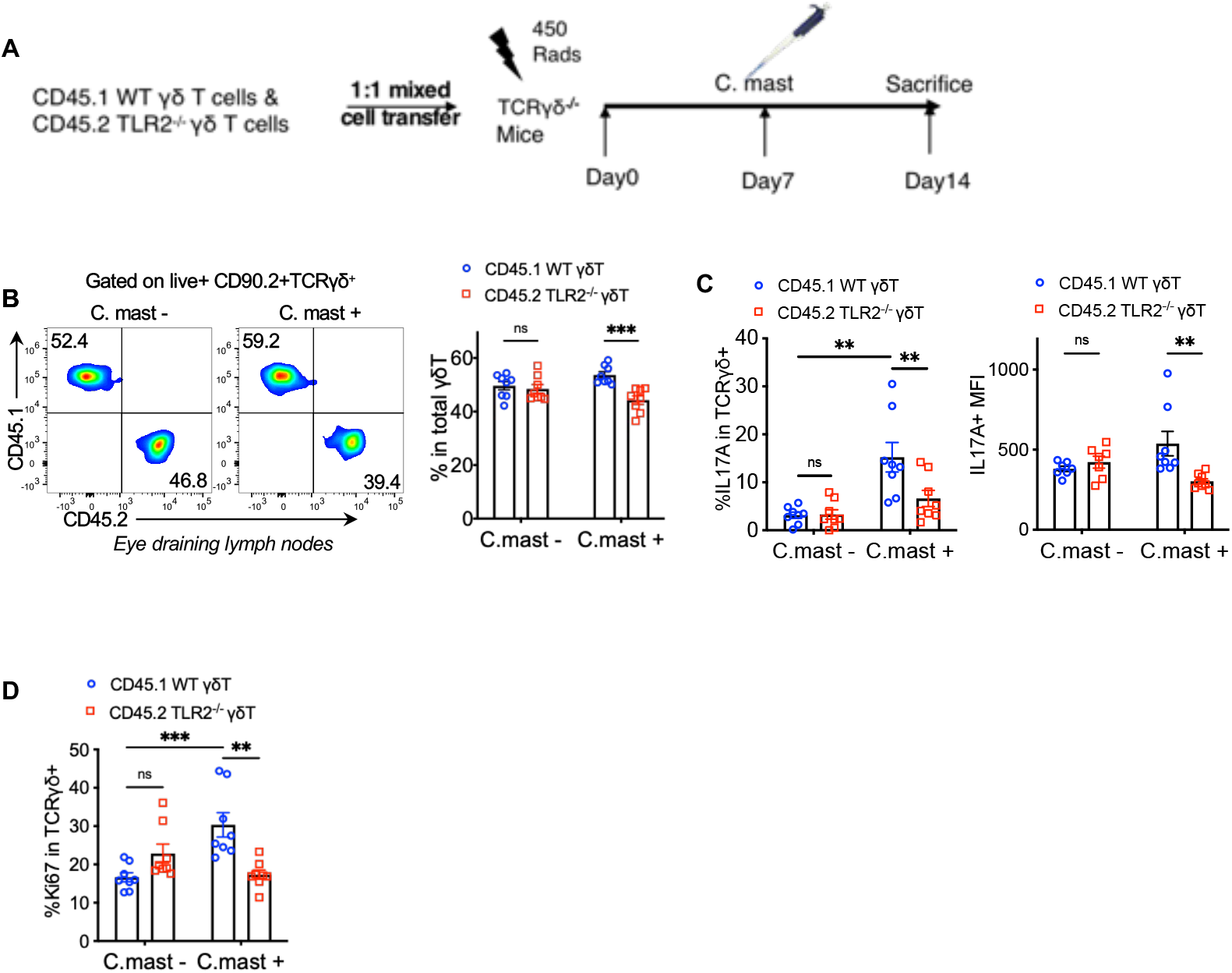
TLR2 deficiency in γδ T cells blunts IL-17A and proliferative responses *in vivo*. A 1:1 mix of CD45 allotype-marked WT and TLR2^−/−^ γδ T cells was infused into TCRδ^−/−^ mice. Cells were collected after 2 weeks for *ex vivo* analysis after PMA/Ionomycin stimulation. **(A)**Schematic representation of the γδ T cell adoptive transfer experiment. **(B)**Representative FACS plots and bar plots showing the ratio of CD45.1^+^ WT γδ T cells and CD45.2^+^ TLR2^−/−^ γδ T cells in the eye-draining lymph nodes from host mice 2 weeks after adoptive transfer with/without *C. mast* inoculation. N= 8. **(C)**The percentage of IL-17A+ cells and IL-17A expression (MFI) of WT and TLR2^−/−^ γδ T cells in the eye-draining lymph nodes from host mice after adoptive transfer with/without *C. mast* inoculation. N= 8. **(D)**The percentage of Ki67+ cells from WT and TLR2^−/−^ γδ T cells from host mice with/without *C. mast* inoculation. (B-D) N=8. Data were combined from 4 experiments. Bars represent mean ± SEM with *P<0.05, **P<0.01, ***P <0.001. Statistical significance was determined by two-way ANOVA (B-E).

### TLR2^−/−^ Vγ6 cells exhibit a more profoundly impaired response to *C. mast than* Vγ4 cells

The function of γδ Τ cells in terms of cytokine production is closely linked to their usage of specific TCR repertoires. γδ T cell subsets expressing Vγ1, Vγ4, and Vγ6 (Heilig and Tonegawa’s nomenclature) produce IL-17A^24-26^. In our previous study, we identified IL-17A-producing Vγ4 cells at the ocular surface of *C. mast-*colonized mice^12^. We had also observed a population of Vγ4∫ γδ T cells that produced IL-17, but did not characterize their TCR or their requirements for stimulation. In the course of the current experiments we noted that ocular *C. mast* association increased the number of the Vγ1^−^Vγ4^−^ Τ cells that produced IL-17A in the conjunctiva (Fig. 4A). Using a TCR-specific antibody that recently became available (1C10-1F7), we confirmed that conjunctival Vγ4^**–**^ cells expressed the Vγ6 TCR^28^ (Fig. S2). Analysis of eye DLNs revealed the presence of IL-17A^+^ Vγ6 cells even in *C. mast*-naïve mice, but the number and the IL-17A-producing capacity of these cells increased considerably after *C. mast* inoculation (Fig. 4B-C). To connect the Vγ6 cells functionally to the ocular IL-17A response to *C. mast*, we compared the response to *C. mast* of ΤCRγ6 knockout (KO) mice (generouds gift from Y. Belkaid, NIAID, NIH) to their WT littermates. ΤCRγ6 KO mice possess a 23-base-pair deletion in the *TcrgV6* gene, resulting in a systemic lack of Vγ6 cells. Vγ6 cells account for 50% to 60% of conjunctival γδ Τ cells in WT mice, whereas they are undetectable in eye-draining lymph nodes and conjunctiva of ΤCRγ6 KO mice (Fig. S3). Lack of Vγ6 cells did not cause a compensatory increase in Vγ4 numbers in the conjunctiva, which resulted in a reduced total number of conjunctival γδ T cells (Fig. 4D, Fig. S4), as well as reduced neutrophil recruitment (a surrogate marker of IL-17A availability) (Fig. 4E). Importantly, following *C. mast* association, TCRγ6 KO mice challenged with ocular *Pseudomonas aeruginosa* (*P. aeruginosa*) demonstrated a compromised host defense against *P. aeruginosa*, as evidenced by an increased bacterial burden in eyes of TCRγ6 KO(Fig. 4E). Together, these data support that the presence of Vγ6 cells are functionally required for elicitation of a full ocular protective response by *C. mast*.

**Figure 4.**
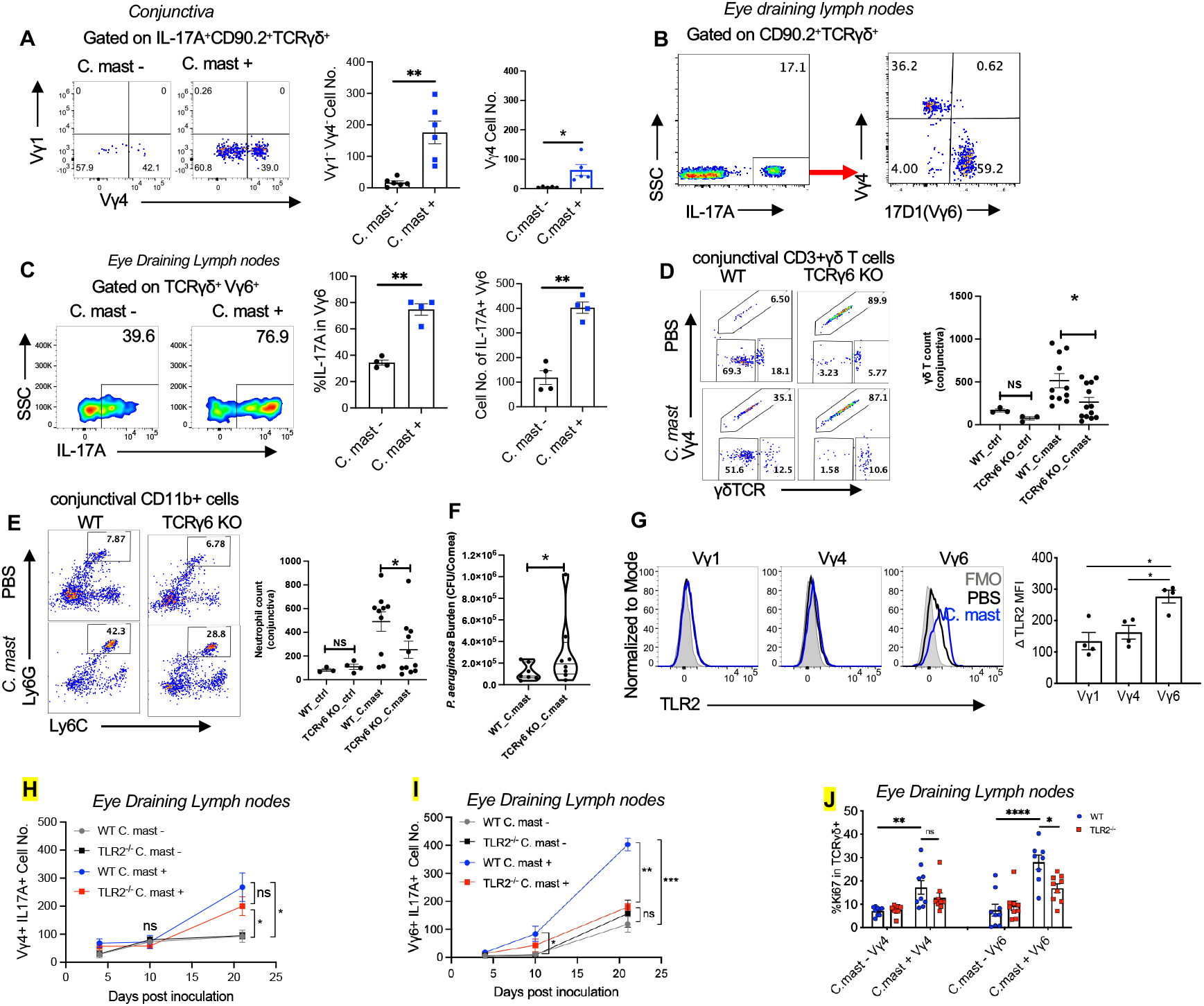
TLR2^−/−^ Vγ6 cells exhibit a more profoundly impaired response to *C. mast than* Vγ4 cells. *C. mast* (CFU =10^8^) or PBS was instilled into each eye of C57BL/6J wild-type mice**(A-C)** every 3 days for a total of 3 times. **(A)**Representative FACS plots and bar graphs representing the number of IL-17A^+^ Vγ1^−^Vγ4^−^ γδ T and IL-17A^+^ Vγ4^+^ γδ T cells in the conjunctiva of wild-type mice with/without *C. mast* inoculation. N= 6. Data were combined from 2 experiments. **(B)**Representative FACS plots showing the percentage of Vγ4^+^, and Vγ6^+^ cells among IL-17A producing γδ T cells of eye-draining lymph nodes with *C. mast* inoculation. The results were representative of 2 independent experiments. **(C)**The percentage and the number of IL-17^+^ Vγ6 cells in eye-draining lymph nodes with or without *C. mast* inoculation. N= 4. **(D-F)**WT and TCRγ6 KO mice were inoculated with either *C. mast* (5 ×10^5^ CFU) or PBS every other day for three inoculations. 7 days after final inoculation, the conjunctiva and draining LNs were harvested. **(D)**Representative FACS plots showing the percentage of Vγ4, Vγ4-γδ TCR^low^ and Vγ4-γδ TCR^high^ γδ T cells in the conjunctiva of indicated mice with/without *C. mast* inoculation. Dot plots showing the number of conjunctival γδ T cells. Each dot represents one animal. **(E)**Representative FACS plots and bar graphs showing the percentage and cell number of conjunctival neutrophils in WT and TCRγ6 KO mice with/without *C. mast* inoculation. Each dot represents one animal. **(F)**7 days after the final *C. mast* association, WT and TCRγ6 KO mice were challenged with *P. aeruginosa (*1 ×10^5^ CFU/per eye). Twenty-four hours later, the affected eyes were harvested and homogenized in PBS. Bacterial load was determined by serial dilution. (N=8 per group). The results were representative of 2 independent experiments. **(G)**Representative FACS plots representing TLR2 expression in the indicated γδ Τ cell subset and bar graphs depicting normalized TLR2 mean fluorescein intensity (denoted as Δ MFI) of the indicated γδ subsets in eye draining lymph nodes. N=4. (Δ MFI = averaged MFI of TLR2 in *C. mast*^+^ group – Averaged MFI of TLR2 in *C. mast*^*–*^ group) **(H-I)** Kinetic experiment showing cell numbers of IL17A^+^ Vγ4^+^ **(E)** and IL17A^+^ Vγ6^+^ **(F)** from WT and TLR2^−/−^ mice with or without *C. mast* inoculation on the indicated days. N = 4. **(J)** Bar plot showing Ki-67 expression of γδ T cells in Vγ4 and Vγ6 subsets in WT and TLR2^−/−^ mice with/without *C. mast* inoculation. N=17. Combined data from 3 experiments. Bars represent mean ± SEM with *P<0.05, **P<0.01, ***P <0.001, ****P≮0.0001. Statistical significance was determined by Welch’s t-test (A,C,F) or two-way ANOVA (D-E, G-J).

Interestingly, the expression of TLR2 became significantly elevated on Vγ6 γδ T cells after *C. mast* colonization compared to other populations of γδ T cells (Fig. 4G). The elevated expression of TLR2 on Vγ6 γδ T cells raised the possibility that TLR2 may have divergent roles in Vγ4 and Vγ6 IL-17A-producing γδ T (γδ T17) cells. We therefore examined the kinetics of IL-17A production by Vγ4 and Vγ6 T cells in the DLNs in response to *C. mast*. The response of Vγ6 cells occurred earlier and was stronger than that of Vγ4 cells, occurring by day 10 in WT mice (Fig. 4H-I) and TLR2 deficiency compromised IL-17A production and proliferation (Ki67) of Vγ6 T cells to a greater extent than that of Vγ4 T cells (Fig. 4H-J). Together, these data lead to the conclusion that Vγ6 T cells are more dependent on TLR2 signaling than are Vγ4 T cells for their responses to the eye-colonizing commensal.

### TLR2 signals regulate the transcriptomic and epigenomic production of IL-17A in Vγ6 cells

To study the basis for the requirement of TLR2 signals in the Vγ6 T subset, we performed RNA-seq on Vγ6 cells sorted from mice prior to and after *C. mast* inoculation (Fig. S5A). The principal component analysis (PCA) plot of the transcriptome revealed distinct clusters formed by WT and TLR2^**−/−**^ Vγ6 γδ T cells (Fig. 5A). Interestingly, both C. mast-negative groups show broad variability; presence of C. mast appears to dramatically focus the response and reduce that variability. Consistent with observations presented in Figure 4, the Vγ6 Τ cells from TLR2^−/−^ mice displayed lower expression of *Il17a* -related genes, such as *Il17a, Il17f, Il17ra, Rorc*, and *Ccr6* (Fig. 5B), confirming that the reduced production of IL-17A protein in TLR2^**−/−**^ Vγ6 γδ T cells is regulated at the transcriptional level. We also investigated the chromatin accessibility of *Il17a* loci in WT and TLR2^**−/−**^ Vγ6 cells by ATAC-seq. Two open chromatin regions (OCRs) were called in *Il17a* loci of Vγ6 cells (Fig. 5C). Based on their genomic location and literature, we parsed one OCR in the promoter region of *Il17a* gene and the other OCR located at the conserved cis-regulatory element 2 (CNS2) region approximately 5500bp upstream of *Il17a* transcriptional start site^29,30^. Comparing chromatin accessibility in *Il17a* loci of other IL-17A-producing cells, such as Vγ4 and ILC3 cells, as reported by ImmGen Consortium, we observed a prominent peak at the CNS2 region of their *Il17a* loci, confirming that this region serves as a conserved cis-regulatory element of *Il17a* loci in these IL-17A producing immune cells (Fig. S5B). Vγ6 cells from *C. mast-*associated WT mice displayed significantly higher accessibility in *the Il17a* CNS2 region than those from *C. mast-*associated TLR2^**−/−**^ mice (Fig. 5C). This finding corroborates that TLR2 signals regulate IL-17A production in Vγ6 cells at the transcriptomic and epigenomic levels.

**Figure 5.**
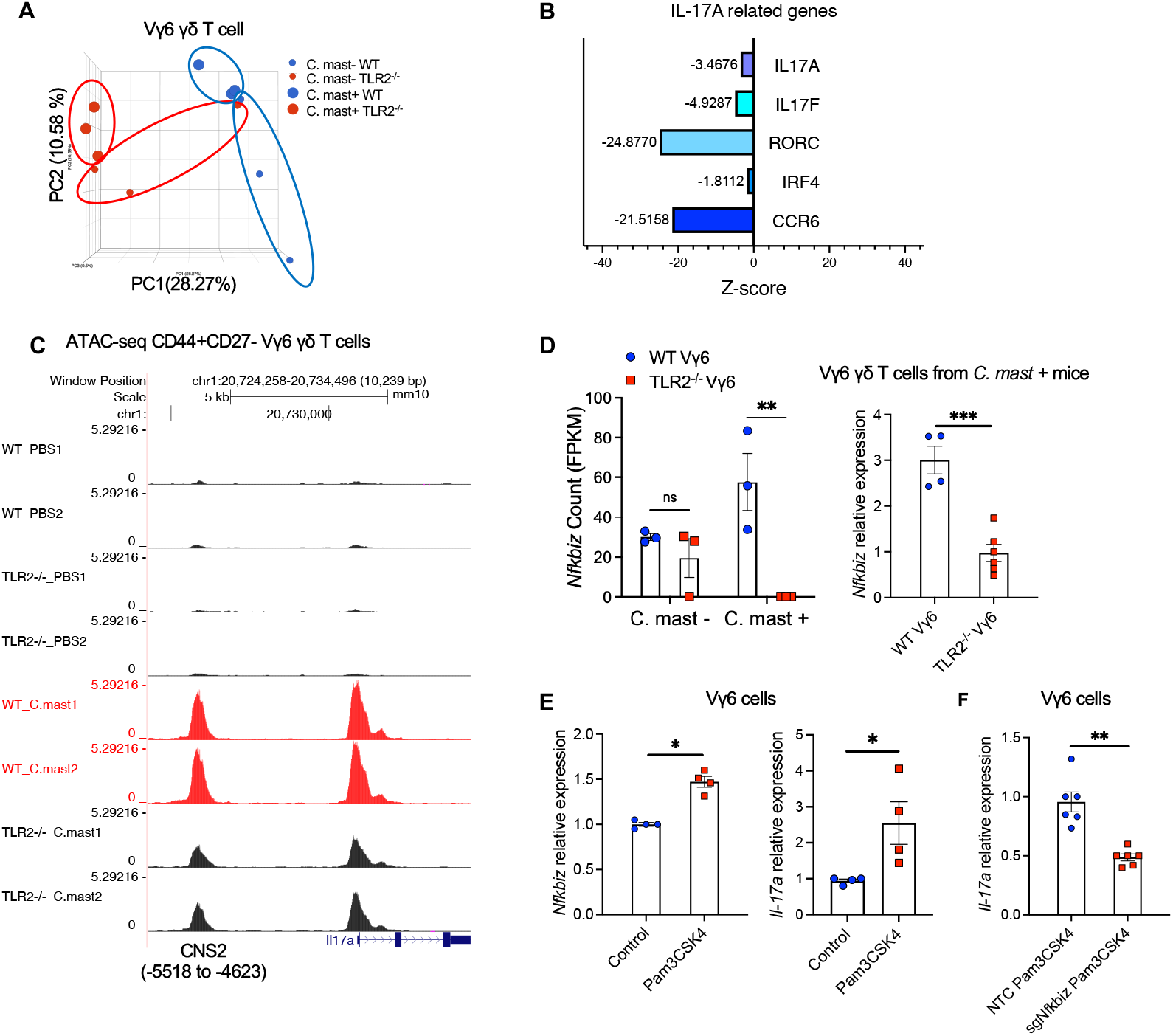
Compromised IL-17A response in TLR2-deficient Vγ6 cells is due to reduced expression of IκBζ. TLR2^−/−^ or WT mice were associated with *C. mast* on their ocular surface. **(A)**RNA-seq was performed on Vγ6 T cells isolated from the indicated mice. The PCA plot shows 4 groups: cells from naïve WT or TLR2^−/−^ mice, and cells from *C. mast*^+^ WT or TLR2^−/−^ mice. N= 3 per group. **(B)**Z scores of selected IL-17 pathway-related genes in WT and TLR2^−/−^ Vγ6 cells from *C. mast*^+^ mice. Blue bars represent genes downregulated in TLR2^−/−^ Vγ6 cells. Z scores = (mean gene counts in TLR2^−/−^ *C. mast*^+^ group – mean gene counts in WT *C. mast*^+^ group)/ SD of corresponding WT *C. mast*+ group’s counts. **(C)**ATAC-seq data showing traces, to the same scale, for Il17a genomic region of CD27-CD44^high^ Vγ6 from C. mast^+^ WT and TLR2^−/−^ mice. N= 2 per group. **(D)**Bar plots show the reads of *nfkbiz* (encodes IκBζ) from RNA-seq of Vγ6 cells. N= 3 per group. Vγ1^−^Vγ4^−^ cells(Vγ6)were sorted from WT and TLR2^−/−^ mice after *C* .*mast* inoculation. The gene expression of *nfkbiz* in sorted Vγ6 cells was quantified by real-time PCR and normalized to β-actin. N= 4-6 per group. **(E)**Vγ6 cells were stimulated with IL-7 alone (Control) or IL-7 plus TLR2 ligand Pam3CSK4 for 2 days. The expression of *Nfkbiz* and *Il17a* was quantified by real-time PCR and normalized to β-actin. Data were combined from 2 experiments. N=4. **(F)**Vγ6 cells were transfected with pooled CRISPR ribonucleoprotein complexes, targeting different regions of Nfkbiz (sg1 and sg2) or a non-targeting control (NTC) that does not target the mouse reference genomes. Three days later, CRISPR-edited samples stimulated with PAM3CSK4 for one day were evaluated for Il-17a expression by real-time PCR. Data were combined from 2 experiments. N=6. Bars represent mean ± SEM. Significance was determined by Welch’s t-test (D right box-F) or two-way ANOVA (D left box).

To identify TLR2-mediated transcription factors (TFs) that influence accessibility of the *Il17a* CNS2 region, we screened TFs known to bind to the *Il17a* CNS2 region and compared their expression between WT and TLR2 deficient Vγ6 cells detected by RNAseq. Of particular interest to us was IκBζ (encoded by gene *nfkbiz)* because of its capacity to bind the *il17a* CNS2 locus in Th17 cells^31^. Furthermore, its transcript levels correlated positively with TLR2 transcript expression quantified by RNAseq and real-time PCR (Fig. 5D).

To examine whether TLR2 activation increases the expression of IκBζ, which in turn promotes transcription of *Il17a*, we sorted Vγ6 cells from Vγ6 TCR-transgenic mice^32^, treated them with the TLR2 ligand Pam3CSK4 in vitro, and measured the gene expression of IκBζ and IL-17A. As expected, Vγ6 cells after Pam3CSK4 stimulation expressed more transcripts of both *nfkbiz* and *Il17a* compared to stimulation with IL-7 (Fig. 5E). Next, we generated IκBζ-deficient Vγ6 cells using a Cas9/gRNA ribonucleoprotein (RNP) transfection that targeted *nfkbiz* (Fig. S5C). Compared to Vγ6 cells transfected with a non-targeting control, IκBζ-deficient Vγ6 cells had diminished *Il17a* transcription (Fig. 5F). These findings indicate that TLR2 in Vγ6 cells promotes IL-17A production at the transcriptomic and epigenomic levels, at least in part through the involvement of IκBζ.

### TLR2-deficient Vγ6 T cells show impaired fatty acid oxidation

Ingenuity Pathway Analysis (IPA) of differentially expressed genes in *C. mast-*associated mice revealed that WT Vγ6 T cells were enriched in the Oxidative Phosphorylation (OXPHOS) pathway (Fig. 6A). OXPHOS takes place in mitochondria and generates ATP through oxidation of nutrients to meet cellular energy demands. We assessed the mitochondrial mass in Vγ6 T cells using MitoTracker Green staining. Both WT and TLR2^−/−^ Vγ6 cells displayed a similar increase in mitochondrial mass upon *C. mast* association, indicating TLR2 deficiency did not affect mitochondrial mass (Fig. S6A). The percentage of MitoSOX (mitochondria superoxide indicator)-positive cells was also comparable between TLR2^−/−^ Vγ6 cells and WT Vγ6 cells (Fig. S6B). In contrast, MitoTracker™ Red CMXros staining, which measures mitochondrial membrane potential (ΔΨm), was reduced in *C. mast-*associated TLR2^−/−^ Vγ6 cells (Fig. 6B), implying impaired mitochondrial activity. This was supported by a reduced ATP production in TLR2^−/−^ Vγ6 cells compared to WT Vγ6 cells (Fig. 6C).

**Figure 6.**
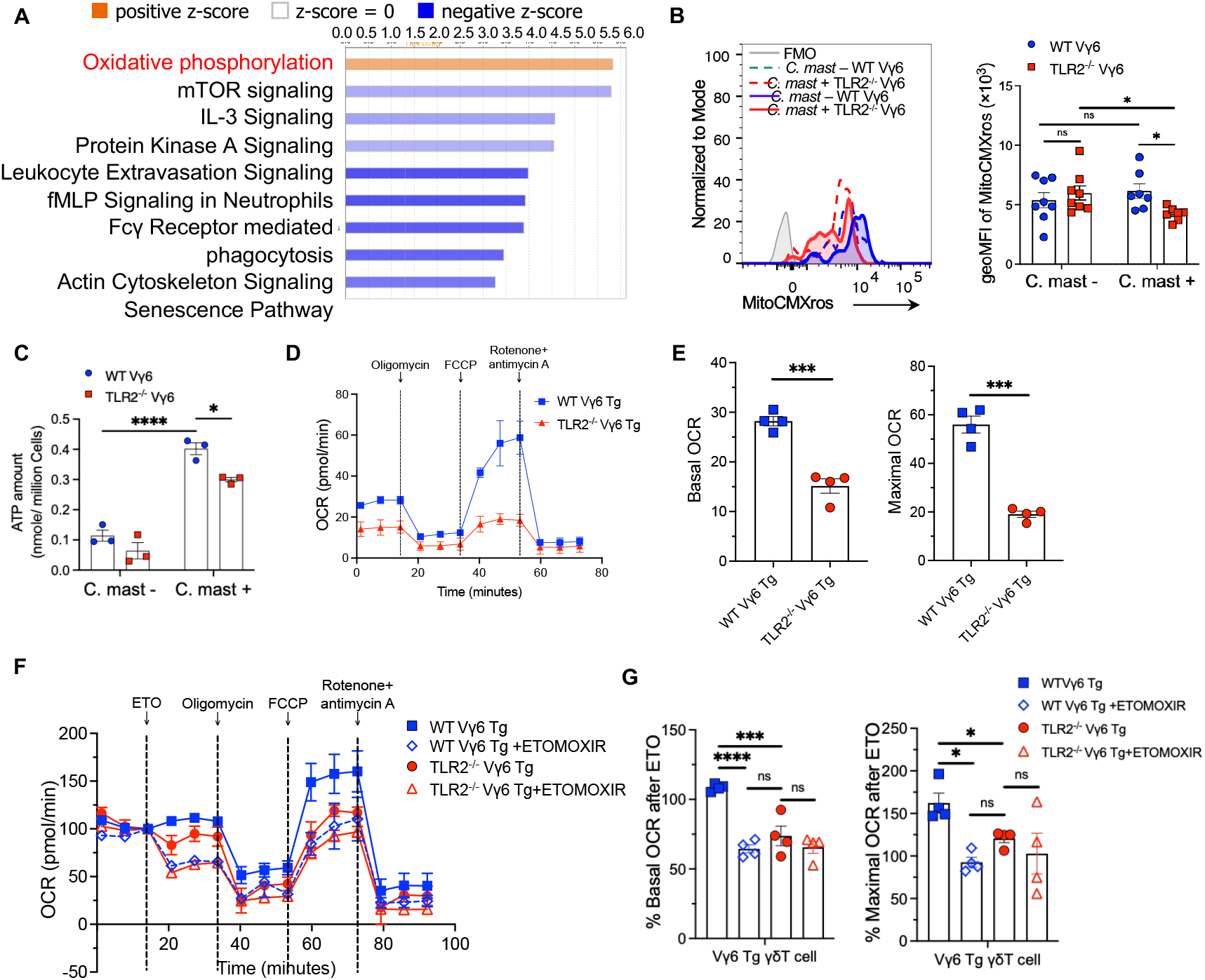
TLR2 deficient Vγ6 T cells show impaired FAO. γδ cells were collected from secondary lymphoid tissues of TLR2 ^−/−^ or WT mice and subjected to transcriptomic or Seahorse metabolic profiling. **(A)**Differentially expressed genes between *C. mast*^+^ WT Vγ6 T cells vs *C. mast*^+^ TLR2^−/−^ Vγ6 T cells were subjected to Ingenuity Pathway Analysis. Orange bars represent pathways enriched in *C. mast*^+^ WT Vγ6 T cells. Blue bars represent pathways decreased in *C. mast*^+^ WT Vγ6 T cells. **(B)**Histogram and bar graphs showing the expression of mitochondrial membrane potential (MitoCMXros) in Vγ6 cells from WT and TLR2^−/−^ mice with/without *C. mast* association. N= 8. **(C)**Measurement of ATP levels in PBS and *C. mast-treated* Vγ6 cells from WT or TLR2^−/−^ mice. N=3. **(D-E)** OCR (oxygen consumption rate) assessment of WT and TLR2^−/−^ Vγ6 T cells from *C. mast*^*+*^ Vγ6 transgenic (Tg) mice. Cells were sorted from eye-draining lymph nodes and treated with PMA + Ionomycin for 2h before the OCR measurement. Combined data from 2 independent experiments. The OCR was measured following treatment with Oligomycin (2.5 μM), FCCP(2.5 μM), and Rotenone + antimycin A (1μM). The percentages of basal and maximal OCR after inhibitor injection were calculated and are displayed in panel **E**. **(F-G)** A seahorse substrate oxidation stress test was performed in *C. mast*+ WT and TLR2^−/−^ Vγ6 cells using the inhibitor ETOMOXIR. The percentages of basal and maximal OCR after inhibitor injection were calculated and are displayed in panel **G**. N= 4. Combined data from 2 independent experiments. Significance was determined by the two-way ANOVA in (B-C, G) or Welch’s t-test (E).

T Cells can generate ATP through the OXPHOS and/or glycolysis pathways. To determine which ATP production pathway is affected in TLR2^−/−^ Vγ6 cells, we used metabolic flux analysis. To obtain sufficient TLR2^**−/−**^ Vγ6 cells, we crossed Vγ6 TCR-transgenic mice ^32^ onto a TLR2^−/−^ background (denoted as WT Vγ6 Tg or TLR2^−/−^ Vγ6 Tg, respectively). First, we wanted to verify that TLR2^−/−^ Vγ6 Tg (double mutant) cells recapitulated the phenotype of Vγ6 cells from TLR2^−/−^ mice with a conventional TCR repertoire. Stimulation with *C. mast* in the presence of WT DC confirmed that TLR2^−/−^ Vγ6 Tg cells had similarly deficient IL-17A and ATP production to TLR2^−/−^ Vγ6 cells from mice with a WT TCR repertoire (Fig. S6C). Next, we performed metabolic flux analysis comparing WT Vγ6 and TLR2^−/−^ Vγ6 Tg cells using a Seahorse assay. The mitochondrial stress test revealed a significant decrease in basic and maximal oxygen consumption rate (OCR) in the TLR2^−/−^ Vγ6 Tg cells compared to WT Vγ6 Tg cells (Fig. 6D-E). The cell-intrinsic role of TLR2 on OXPHOS was confirmed by direct Pam3CSK4 stimulation of Vγ6 cells *in vitro* (Fig. S6D). In contrast, the glycolytic proton efflux rate (glycoPER), which measures glycolysis, showed no differences between the two groups (Fig. S6E). Taken together, these findings indicated that, glycolysis was unaffected, but TLR2^−/−^ Vγ6 cells exhibited impaired OXPHOS, leading to insufficient ATP production in these cells.

ATP production in mitochondria mainly comes from three substrates: glucose, long-chain fatty acid (LCFA), and glutamine. To determine the dominant substrate(s) in the energy metabolism of Vγ6 cells, we performed the Substrate Oxidation Stress Test on Vγ6 Tg cells from TLR2-sufficient Vγ6 Tg mice. Only etomoxir, an inhibitor of the LCFA oxidation pathway, significantly decreased both basic and maximal OCR in Vγ6 Tg cells (Fig. S6F), demonstrating that γδΤ17 cells utilize primarily fatty acid oxidation for energy production. To test whether TLR2 deficiency impairs the LCFA pathway in Vγ6 cells, we performed a Substrate Oxidation Stress Test on Vγ6 cells from WT Vγ6 Τg and TLR2^−/−^ Vγ6 Τg mice inoculated with *C. mast*. Vγ6 cells from TLR2^−/−^ Vγ6 Τg mice displayed a reduced basic and maximal OCR compared to WT Vγ6 Τg mice after etomoxir treatment (Fig. 6 F-G), consistent with the reduced ATP content in TLR2^−/−^ Vγ6 cells (Fig. 6C). These data confirm that TLR2 signaling supports fatty acid oxidation (FAO) in Vγ6 cells.

### Impaired Cpt1 function underlies the defective IL-17A response of TLR2-deficient Vγ6 cells

To further explore the underlying mechanisms, we studied the impact of TLR2 signaling on fatty acid oxidation (FAO) in Vγ6 γδ Τ cells. ATAC-seq data revealed that *C. mast* association activated an OCR within 2kb upstream of the *Cpt1a* gene, which encodes carnitine palmitoyl transferase 1a, a key enzyme involved in FAO (Fig. 7A). In addition, analysis of ImmGen Consortium data ^33^ identified the same OCR in the *Cpt1a* locus in IL-17A-producing ILC3s from small intestine, which harbors commensal microbiota (Fig. S7). Notably, TLR2 deficiency decreased chromatin accessibility at *Cpt1a* (Fig. 7A). These data suggest that TLR2 signaling can enhance FAO by promoting the transcription of *Cpt1a*, which is also supported by our *in vitro* data showing that Pam3CSK4 stimulation upregulated *Cpt1a* in Vγ6 cells (Fig. 7B).

**Figure 7.**
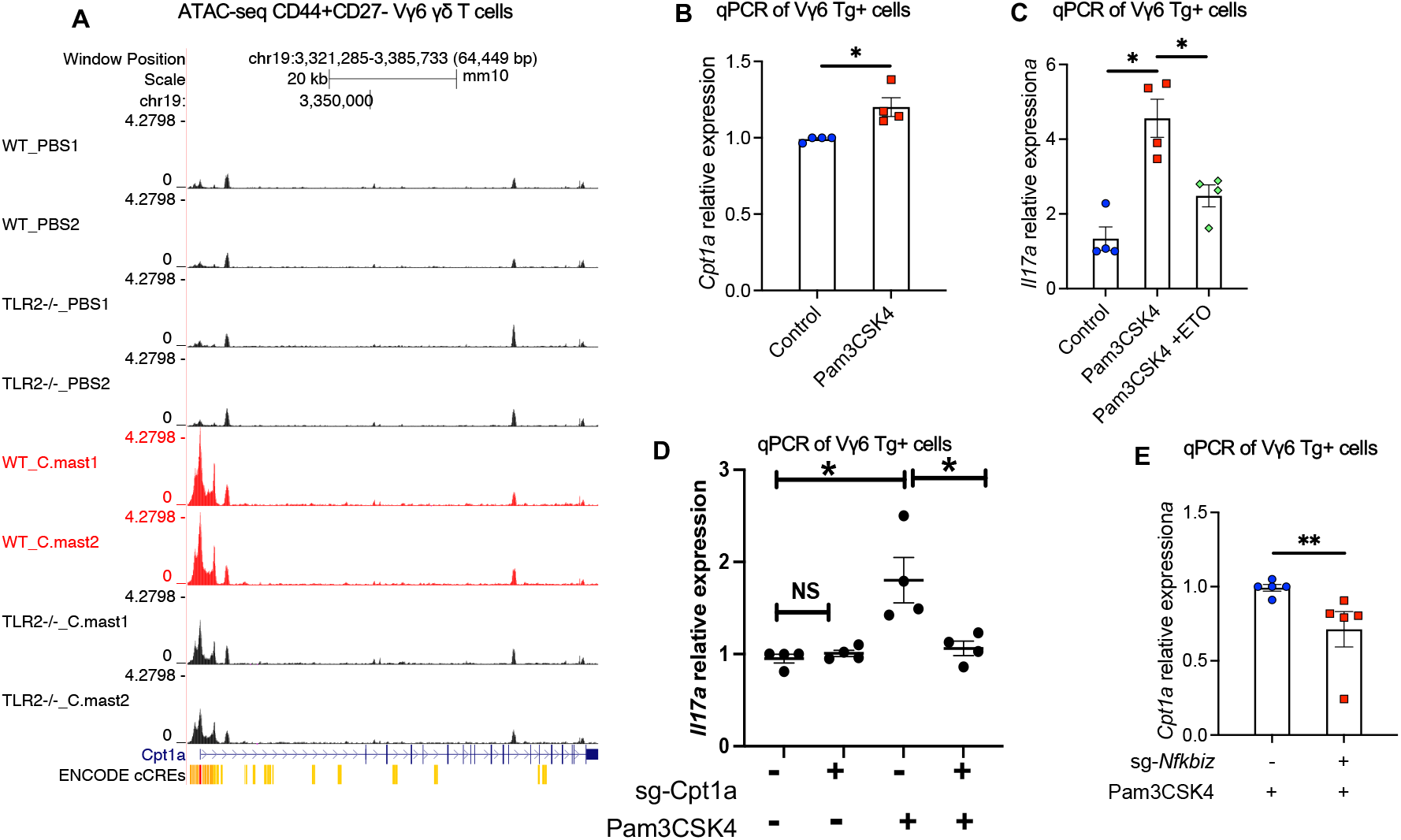
Impaired Cpt1 function underlies the defective IL-17A response of TLR2-deficient Vγ6 T cells. **(A)**UCSC genome browser output depicting ATAC-seq traces for the promoter and enhancer regions (ENCODE cCREs) of *Cpt1a* loci in CD27-CD44^high^ Vγ6 cells from WT and TLR2^−/−^ mice. **(B)**Vγ6 cells were sorted from *C. mast-* Vγ6 Τg mice and stimulated with IL-7 alone (Control) or IL-7 + Pam3CSK4 (Pam3CSK4) for one day. Expression of the *Cpt1a* gene is quantified by real-time PCR and normalized by the β-actin. N=4. **(C)**Vγ6 cells were stimulated with IL-7 alone (Control) or IL-7 plus TLR2 ligand Pam3CSK4 (Pam3CSK4) for 2 days, Etomoxir (40 μM) was added for the final 12 hours. Expression of *Il17a* was quantified by real-time PCR and normalized to β-actin. N=4. **(D)**Vγ6 cells were transfected with pooled CRISPR ribonucleoprotein complexes targeting different regions of Cpt1a, or a non-targeting control (NTC) not targeting mouse reference genomes. Three days later, CRISPR-edited samples were stimulated with Pam3CSK4 for one day and evaluated for *Il17a* expression by real-time PCR. N=4. Each dot in the graph represents a repeated experiment. **(E)**Vγ6 cells were transfected with pooled CRISPR ribonucleoprotein complexes targeting different regions of Nfkbiz, or a non-targeting control (NTC) that did not target mouse reference genomes. Three days later, CRISPR-edited samples were stimulated with Pam3CSK4 for one day and evaluated for *Cpt1a* expression by real-time PCR. N=5. **(B-C,E)** Representative data of 2 independent experiments. Significance was determined by the Mann-Whitney test (B,E) or two-way ANOVA in (C-D).

We next examined directly whether FAO is involved in TLR2-mediated IL-17 response in Vγ6 cells. Treatment with etomoxir at 40 μM, a concentration that inhibits Cpt1a without off-target^34^ effects, blocked the upregulation of *Il17a* induced by TLR2 activation (Fig. 7C). Consistent with this result, Cpt1a-insufficient Vγ6 cells that were generated by a Cas9/gRNA ribonucleoprotein (RNP) transfection that targeted *Cpt1a* (Fig. S8), lost the ability to upregulate *Il17a* in response to TLR2 activation (Fig. 7D). Notably, IκBζ, which in our hands upregulates *Il17a* in Vγ6 cells in response to TLR2 signaling, had also been reported to upregulate *Cpt1a* in fibroblastic reticular cells^35^. In line with this, IκBζ-insufficient Vγ6 cells that we generated by Cas9/Crispr targeting, displayed reduced *Cpt1a* transcription (Fig. 7E). In the aggregate, our data support a sequence of events where, in response to endogenous TLR2 signaling in Vγ6 cells, increased expression of IκBζ enhances transcription of Cpt1a, thus favoring fatty acid oxidation and providing metabolic support for IL-17A production.

## DISCUSSION

Our earlier studies demonstrated that the ocular commensal *C. mast* elicits expansion and IL-17A production by γδ Τ cells that provide local defense from pathogens at the ocular surface mucosal barrier. The current study explores mechanistically how these γδ T cells sense and respond to *C. mast*. We demonstrate that *C. mast* association activates TLR2 signals on both DCs and γδ Τ cells. While engagement of TLR2 on DC triggered the IL-1β essential to elicit γδ T cell IL-17 production, intrinsic TLR2 signaling in γδ Τ cells was also required. Our data reveal that intrinsic TLR2 signaling in γδ17 T cells acts in a subset-specific manner to enhance transcription of genes related to IL-17A production and activates the metabolic reprogramming to fatty acid oxidation that is needed to support IL-17A production.

Microorganisms activate TLR2 signaling in antigen-presenting cells and Τ cells (both αβ and γδ Τ cells) for IL-17A responses^17^. In our investigation of *C. mast*, we were able to dissect the contribution of TLR2 expression in DCs from that of γδ Τ cells. Specifically, TLR2 activation in DCs by *C. mast* induced IL-1β, facilitating the proliferation and IL-17A production in γδ T cells. While both Vγ4 and Vγ6 cells exhibited IL-17A responses to *C. mast*, intrinsic TLR2 signals play a dominant role in Vγ6, but not Vγ4, cells for their proliferation and effector functions. The greater reliance on TLR2 signals for IL-17A responses in Vγ6 cells stimulated by *C. mast* are reminiscent of other reported phenotypes, e.g., in mouse gingiva, where the presence of Vγ6, but not Vγ4, cells is dependent on the presence of commensals^25,36^. Based on our findings, we speculate that activation of innate receptors such as TLR2, by these commensals, may explain persistence in the gingival Vγ6, cells but not for Vγ4 cells. These phenotypes may perhaps be connected to the inherited innate-like behavior of Vγ6 cells towards *C. mast* and apparently also other commensals^37,38^. Vγ4 cells, on the other hand, are later-stage responders, which may be explained by their need to receive TCR stimulation in addition to signals provided by DCs^12^.

TLR2 signals on Vγ6 cells target transcription factors in IL-17A regulation. We observed that TLR2 deficient Vγ6 cells displayed a decrease in the transcriptional expression of IκBζ. This transcription factor was found to upregulate *Il17a* in αβ Th17 cells^31^. However, the need for IκBζ in IL-17A production by γδT17 cells has not been reported, and in view of the very different developmental and functional niches of these lineages, was not a given. In our hands, an *in vitro* knockout of IκBζ reduced *Il17a* transcript levels in Vγ6 cells, suggesting a supportive role of IκBζ in Vγ6 cells’ IL-17A pathways. In addition to transcriptional regulation, we observed the epigenomic regulation mediated by TLR2 signaling at *Il17a* locus in Vγ6 cells, especially at the promoter and CNS2 region. The transcription factor IκBζ binds to the *Il17a* CNS2 region in αβ Th17 cells, facilitating IL-17 transcription^31^. Although our study did not directly examine the binding of IκBζ to the *Il17a* regulatory region in Vγ6 cells, it stands to reason that lower binding of IκBζ to *Il17a* CNS2 region would contribute to the decreased *Il17a* transcripts in TLR2-deficienct Vγ6 cells. Notably, IκBζ does not act alone; it cooperates with Rorc to stimulate *Il17a* transcription in Th17 cells^31,39^. Our RNA-sequencing data revealed a positive association between Rorc and TLR2 sufficiency, which supports the notion that Rorc upregulates Il17a through TLR2 signaling in Vγ6 cells. These findings suggest that TLR2 signaling utilizes multiple transcription factors that work together to regulate the transcriptional and epigenomic IL-17A program in Vγ6 cells.

Recent data made it clear that cell metabolism critically influences the functionality of immune cells. Cell metabolism is largely dictated by energy availability in the form of glucose, fatty acids, or other sources. However, other factors can impact cellular energy profiles. Here, we show that *C. mast*, through TLR2, can activate the OXPHOS pathway in γδ T cells, which then supports the production of IL-17A at the ocular surface. We posit that this finding parallels the situation found in the intestine, where colonizing segmented filamentous bacteria (SFB) can stimulate lipid metabolism in CD4^+^ Th17 cells^40^. Similarly, type 3 innate lymphoid cells (ILC3s) produce IL-17A in a manner that correlates with increased lipid metabolism^41^. These results, together with observations that germ-free mice display impaired host metabolism and IL-17 responses at intestinal mucosal sites^18,42,43^, suggest a shared metabolic regulation network across various mucosal surfaces that depends on the colonization of microbes at specific mucosal sites.

TLR2 signaling, functioning as a sensor of commensal bacteria, is involved in commensal-mediated metabolic adaptation in γδ Τ17 cells. While previous studies have reported the impact of TLR2 signals on proliferation, survival, and cytokine production in various T cell populations, our findings expand the understanding of TLR2 signaling in the metabolic regulation of γδΤ17 cells. TLR2-activated FAO can increase expression of the *Il17a* gene. Mechanistically, TLR2 signaling can upregulate the expression of *Cpt1a* in Vγ6 cells, which correlates with enhanced FAO capacity. Notably, IκBζ, which increases *Il17a* transcription, also enhances the expression of *Cpt1a* in Vγ6 cells. These data suggest a possible feed-forward mechanism where factors downstream of TLR2 signaling can epigenetically influence the future identity and metabolic profile of the cell. The mechanism(s) through which IkBζ promotes Cpt1a expression require(s) further investigation.

In summary, our results highlight the importance of intrinsic TLR2 signaling in γδ Τ cells to produce IL-17A in response to bacterial colonization. TLR2 signals enhance the expression of genes responsible for the IL-17A pathway and fatty acid metabolism in IL-17A-producing γδ Τ cells. Consequently, this study further links IL-17A production in γδ Τ cells with cell metabolism, implicating metabolic intervention as a potential strategy to modify γδ T cell functionality at mucosal and/or other barrier surfaces.

## Supporting information

All supplementary Figures

## STAR METHODS

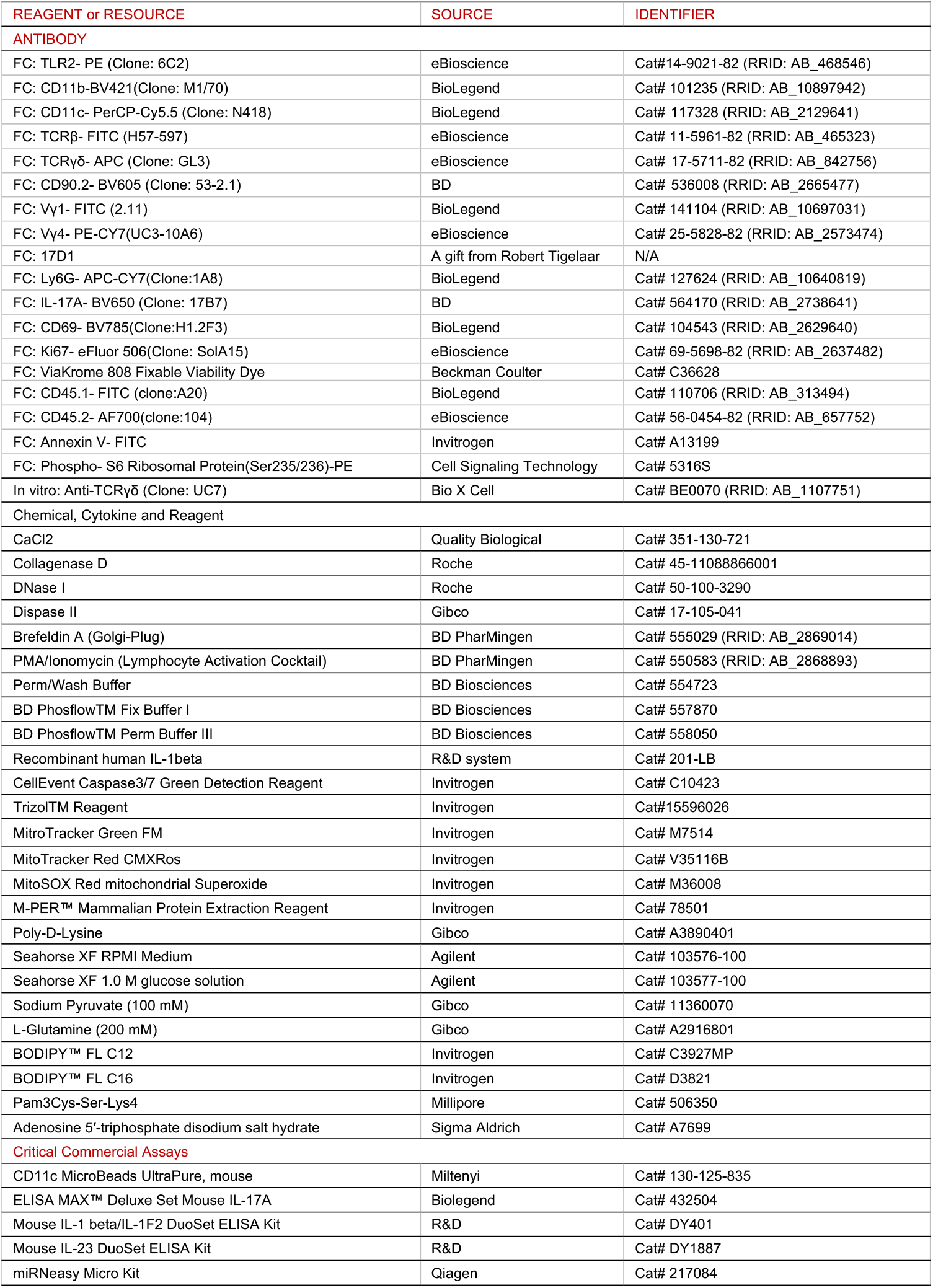
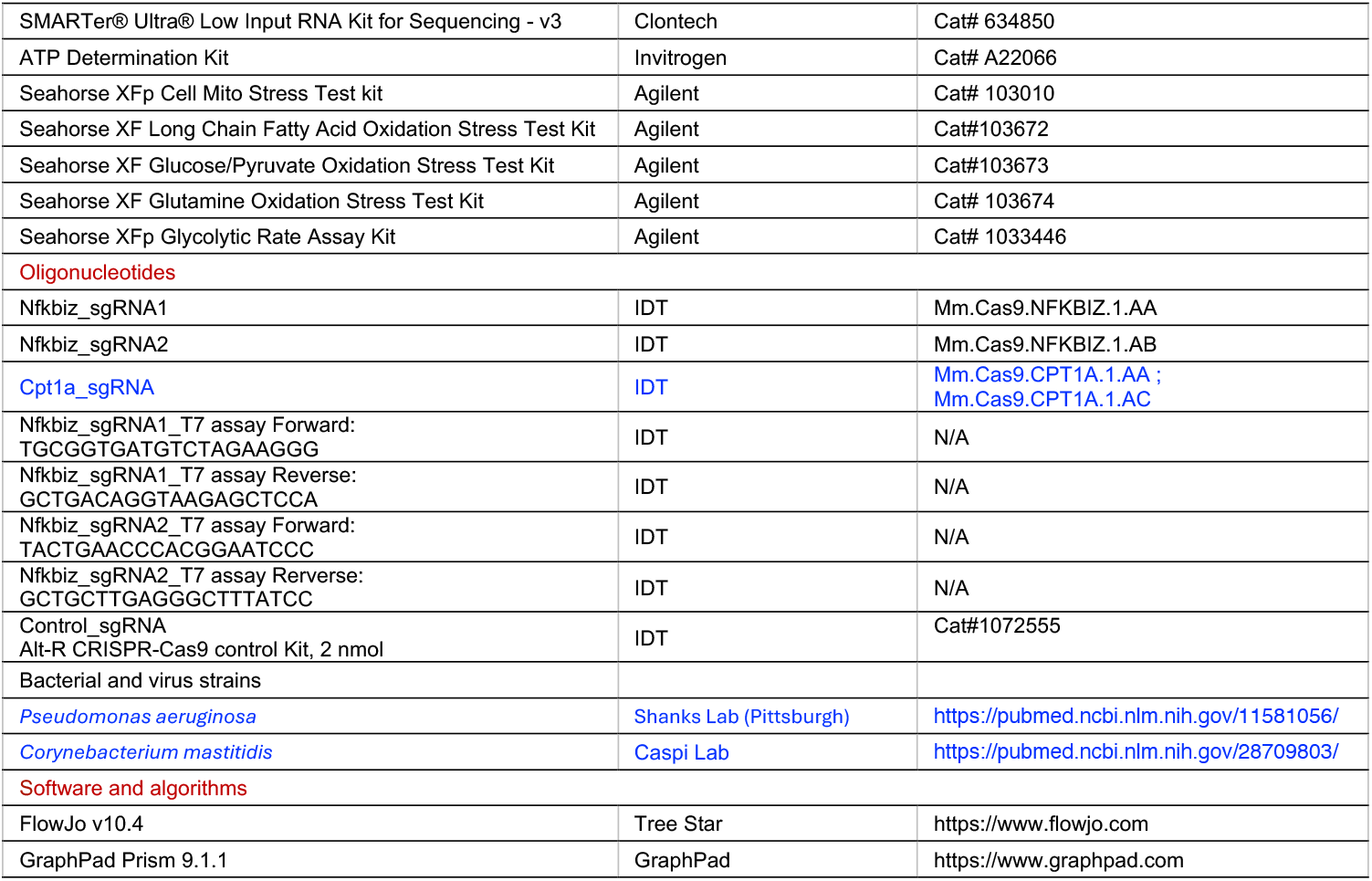
KEY RESOURCES TABLE.

## CONTACT FOR REAGENT AND RESOURCE SHARING

Further information and requests for resources and reagents should be directed to and will be fulfilled by the Lead Contact, Rachel Caspi and/or Xiaoyan Xu

## Data and Code availability

The RNA-seq datasets and the ATAC-seq datasets generated during this study are available at NCBI GEO (<u>GSE236387</u>). The code used for data analysis is shared at https://github.com/NIH-NEI/tlr2-ATAC-seq/

## EXPERIMENTAL MODEL AND SUBJECT DETAILS

### Mice

WT C57BL/6J mice were purchased from Jackson Laboratory. TLR2^−/−^, TCRβ^−/−^, and CD45.1 C57BL/6J mice originally from Jackson were bred and maintained in our animal facility. Vγ6 transgenic mice were generously provided by Paul E. Love; C57BL/6-*Trgv6*^*em1Ybe*^(TCRγ6 KO) mice were provided by Yasmine Belkaid; TLR2^−/−^Vγ6 transgenic mice, and TLR2^−/−^ TCRβ^−/−^ mice were crossed and bred in house. For most in vivo and in vitro animal studies, male and female mice aged 6-10 weeks were used. All mice were housed under NIH SPF conditions. All mice received tear washes and were confirmed to lack ocular surface *C. mast* colonization before the experiment. All mouse studies were performed in full compliance with IACUC-approved protocols (NEI-690) and institutional guidelines.

### Bacteria

*C. mast* was initially isolated from conjunctival homogenates that were plated on TSA + 5% blood agar plates kept at 37°C in anaerobic conditions for 7 days. After original isolation, C. mast was propagated using TSA with 5% blood agar plates at 37°C in an aerobic chamber ^12^. Further, C. mast can be grown in blood heart infusion broth with 1% Tween 80 aerobically with shaking at 37°C.

## METHOD DETAILS

### *C. mast* Inoculation

*C. mast* was applied (10^8^ CFU) to the ocular surface every three days for a total of three inoculations. Three weeks after the final inoculation, 10 μl PBS per eye was used to wash the conjunctiva to confirm colonization.

### Tissue harvest and processing

Mice were euthanized by extensive cardiac perfusion. Two conjunctivae of one mouse were harvested by excising the eyelid and bulbar conjunctiva, combined, minced, and exposed to 0.9 ml of 1 mM CaCl_2_, 2 mg/ml collagenase D, 0.16 mg/ml DNaseI and 0.05 mg/ml Dispase II in HBSS for 45 min at 37°C with shaking. Cervical and submandibular lymph nodes were isolated, minced and exposed to 1mg/ml collagenase D for 30 min at 37°C with shaking. After collagenase treatment, conjunctiva and draining lymph node cells were separately filtered through a 40 μm filter using a 1 cc syringe plunger. Then conjunctival cells were filtered a final time using a Corning Falcon Test Tube with Cell Strainer Snap Cap (Corning 352235).

### CD11c Isolation

Mouse spleen tissue was cut into small pieces in RPMI medium containing 1mg/ml collagenase D. Tissue was incubated at 37°C and placed on a shaker for 1 hour. The single-cell suspension was prepared with a 70um filter and 1cc syringe plunger and was incubated with Miltenyi CD11c Isolation Beads and Fc Blocker for 10 mins on ice. CD11c positive selection was then performed following the manufacturer’s protocol.

### Cell sorting

Singe cell suspensions were prepared from cervical lymph nodes and spleen of WT or TLR2^−/−^ mice. Total γδ T cell sorting: Fc blocker, APC anti-TCRγδ, and BV421 anti-TCRαβ antibody were added to the cell suspension in the FACS buffer and stained for 30 mins at room temperature. For Adoptive transfer: FITC anti-CD45.1, PE anti-TCRγδ, AF700 anti-CD45.2, BV510 anti-CD90.2 were mixed and stained. For RNA-seq cell sorting, FITC anti-Vγ1, PE-CY7 anti-Vγ4, APC anti-TCR γδ, and BV421-TCRαβ, BV510 anti-CD90.2 were mixed and stained cells for 30 mins. For ATAC-seq sorting, FITC anti-Vγ1, PE anti-TCRγδ, PE-CY7 anti-Vγ4, APC anti-CD62L, BV421-TCRαβ, BV510 anti-CD90.2 and BV785 anti-CD44 were mixed and added to cells. Propidium Iodide (PI) was added before each sorting, and cells were isolated by BD FACS Aria cell sorters.

### Preparation for heat-killed *C. mast*

*C. mast* (1×10^6^ CFU) was suspended in 100 μl PBS and heated at 60 °C for 30 mins.

### In vitro γδ T cell and CD11c+ cell co-culture

CD11c^+^ cells (1×10^5^) were co-cultured with TCRγδ^+^ cells (0.2×10^5^) in 200 μl RPMI supplemented with 10% FBS for 72 hours under indicated conditions: 1 μl of heat-killed *C. mast* was added to 200 μl culture medium with 100 ng/mL IL-1β.

### Flow Cytometry

*In vitro* assay: For intracellular staining, Brefeldin A (Golgi Plug) was added for the last 6 hours of 72-hour co-culture. After centrifugation, the supernatant was collected for ELISA assays. Cells were collected for flow cytometry. *In vivo* assay: tissue was homogenized in 2% FBS RPMI and filtered by a 40 μm strainer. The cells were stained following the manufacturer’s protocol. For phospho-signal flow cytometry, we use BD Phosflow Fix Buffer and BD Phosflow Perm/Wash Buffer III. All the procedures followed by BD™ Phosflow Protocols for Mouse Splenocytes or Thymocytes. All samples were acquired with Beckman CytoFLEX LX Flow Cytometer and analyzed with FlowJo software.

### ELISA

Cell supernatant was collected for IL-17A, IL1-β, IL-23 ELISA assays. Biolegend ELISA MAX™ Deluxe Set Mouse IL-17A (Cat# 432504), R&D Mouse IL-1 beta/IL-1F2 DuoSet ELISA (Cat# DY401), R&D Mouse IL-23 DuoSet ELISA Kit (Cat# DY1887) were used according to manufacturer’s protocols.

### Adoptive Transfer Assay

γδ T cells were sorted from CD45.1^+^ WT mice and CD45.2^+^ TLR2^−/−^ mice (0.3 million cells each). Cells were mixed before being transferred to sublethal irradiated (450 rad) TCRγδ^−/−^ mice.

### RNA-seq

PI^-^CD90.2^+^TCRβ^-^TCRγδ^+^Vγ1^+^Vγ4^-^ cells, PI^-^ CD90.2^+^TCRβ^-^TCRγδ^+^Vγ1^-^Vγ4^+^ cells, PI^-^ CD90.2^+^TCRβ^-^TCRγδ^+^Vγ1^-^Vγ4^-^cells were sorted from naïve WT, naïve TLR2^−/−^, *C. mast* inoculated WT and *C. mast* inoculated TLR2^−/−^ mice. RNA was isolated using Qiagen miRNeasy Micro Kit. RNA was run on Agilent 2100 Bioanalyzer and RIN >8.0 was aimed to pass QC. RNA-seq was performed on Illumina Hiseq4000 (150pb, paired) using Clontech RNA Ultra Low Input Library Prep and paired-end sequencing in the NCI sequencing facility. Reads of the samples were trimmed for adapters and low-quality bases using Cutadapt before alignment with the reference genome (Mouse - mm10) and the annotated transcripts using STAR. The mapping statistics were calculated using Picard software. Library complexity is measured in terms of unique fragments in the mapped reads using Picard’s MarkDuplicate utility. In addition, the gene expression quantification analysis was performed for all samples using STAR/RSEM tools.

### ATAC-seq

DAPI^-^CD90.2^+^TCRβ^-^TCRγδ^+^Vγ1^-^Vγ4^-^CD27^-^CD44^hi^ cells were sorted from naïve WT, naïve TLR2^−/−^, *C. mast* inoculated WT, and *C. mast* inoculated TLR2^−/−^ mice. Library preparation and sequencing were done by Novogene based on a previously reported method^44^. Briefly, the nuclei extracted from the cells were quality controlled and suspended in the TruePrep Tagment Enzyme mix and incubated at 37°C for 30 min, followed by PCR amplification and purification with the AMPure beads. The quality-controlled library was quantified with Qubit 2.0, clustered using TruSeq PE Cluster Kit v3-cBot-HS (Illumina) and sequenced on an Illumina Hiseq platform to generate 150 bp paired-end reads. Raw data underwent rigorous quality control through FastqQC and FastP^45^. Alignment was executed with Bowtie 2 ^46^ and peaks were annotated using Genrich^47^. ChiPseeker^48^, and differential expression analysis relied on DESeq2^49^. Homer2 was employed for motif analysis, and the UCSC genome browser ^50^ was the platform for peak visualization.

### Extracellular flux analyses (Seahorse)

Experiments were performed on the Agilent Seahorse XF96 bioanalyzer. Freshly sorted γδ T cells were generated from *C. mast*^+^ WT and TLR2 mice. γδ T cells were stimulated with PMA and Ionomycin for 3 hours before starting the Seahorse assay. Cells were washed with a prepared assay medium. Cells were plated onto XFp microplates (3×10^5^ cells/well) precoated with Poly-D Lysine (Thermo Fisher). The plates were centrifuged to immobilize cells. Cells were rested in assay medium in a non-CO2 incubator for 1hr prior to the assay. Oxidative phosphorylation-associated parameters were determined by Seahorse XFp Cell Mito Stress Test kit (Agilent, #103010) with three injections: (1) 2.5 μM oligomycin, (2) 2.5 μM FCCP, and (3) 1 μM rotenone/antimycin A. For the XF Substrate Oxidation Stress Test kit (Agilent, #103672, #103673, #103674), Oxidative phosphorylation associated parameters were determined with four injections: (1) 2.5 μM oligomycin, (2) 4.0 μM Etomoxir/ 2.0 μM UK5099/ 3.0μM BPTES (3) 2.5 μM FCCP, and (4) 1 μM rotenone/antimycin A. Glycolysis associated parameters were determined by Seahorse XFp Glycolytic Rate Assay Kit (Agilent #1033446) with two injections: (1) 0.5 μM rotenone/ antimycin A; (2) 50 mM 2-DG.

### Mitochondrial ROS measurement

Cells were collected from PBS-treated and *C. mast*-treated WT and TLR2^−/−^ mice. Cells were incubated with a MitoSOX probe (Invitrogen, Cat. # M36008) for 10 min at 37°C. After incubation, cells were washed and stained for other surface markers. Then cells were subjected to flow cytometry analysis.

### ATP level determination

γδ T cells were sorted from *C. mast* inoculated WT and TLR2^−/−^ Vg6 transgenic mice and placed in 96 well, pre-coated with anti-TCRγδ (2μg/ml) the previous night. Cells were harvested 3 days post in vitro activation, washed with RPMI-1640 medium three times, and counted. Cells were rested in 96-well for 3 hours at 37°C. Cells were quickly spun down and lysed in a native lysis buffer (Cell Signaling Technology) for 10 min on ice. Cells were then centrifuged at 2000 g for 10 min and the supernatants were collected. ATP levels were measured according to the manufacturer’s instructions (Invitrogen, #A22066).

### CRISPR mutation of IκBζ and Cpt1a in Vγ6 cells

To mutate IκBζ or Cpt1a in Vγ6 cells, CRISPR-Cas9 ribonucleoprotein (RNP) complexes were delivered into Vγ6 Τg cells using electroporation with the Amaxa Nucleofector system (Lonza). RNP complexes are made up of 150 pmol sgRNA, 10 ug Cas9 nuclease and 20 mM Alt-R Cas9 Electroporation Enhancer (Integrated DNA Technologies, Inc) as described previously (Soyoung A. Oh et al., 2019). sgRNAs were designed using the Crispr guide design software (Integrated DNA Technologies, Inc).

### P. aeruginosa challenge

Five-week-old WT or TCRγ6 KO C57BL/6 mice lacking *C. mast* were inoculated with *C. mast* every other day for three inoculations. 7 days after the final inoculation, mice were challenged with *P. aeruginosa* (1 ×10^5^ CFU/per eye). Twenty-four hours later, the affected eyes were harvested and homogenized in sterile PBS with 5 mm stainless steel beads (Qiagen, Hilden, Germany) in the MP FASTPREP 24 (MPBio, Santa Ana, CA). Homogenized eyes were then serially diluted and plated on LB agar. After overnight culture at 37°C, the number of colonies was quantified.

#### Generation of C57BL/6-*Trgv6*^*em1Ybe*^(TCRγ6 KO) mice

To generate CRISPR guide RNA (gRNA) targeting Tcrg-V6, the oligos 5′-TAGGCAGTCTCACGTCACCTCT-3′ and 5′-AAACAGAGGTGACGTGAGACTGC-3′ (gRNA 1) and 5′-TAGGGGGTCATATGTCATCAAG-3′ and 5′-AAACCTTGATGACATATGACCC-3′ (gRNA 2) were annealed and ligated into the pT7-gRNA vector (Addgene). Linearized DNA was used as a template for *in vitro* transcription (IVT) using the MEGAshortscript T7 Transcription Kit (Thermo Fisher Scientific), which yielded the gRNAs. Cas9 mRNA was generated by using linearized pT3TS-nCas9n (Addgene) as the template for IVT with the mMESSAGE mMACHINE T3 Transcription Kit (Thermo Fisher Scientific). Both the Cas9 mRNA and the gRNAs were purified using the MEGAclear Transcription Clean-Up Kit (Thermo Fisher Scientific) and eluted in RNase-free water. C57BL/6NCr blastocysts were microinjected with Cas9 mRNA (100 ng/mL) and gRNA (100 ng/ mL). Progeny were screened by polymerase chain reaction (PCR) amplification with primers 5′-TGATTTCAGTGCCTCTCCTG-3′ and 5′-CACAGCATTCTTTGGAGACG-3′, followed by Sanger sequencing with the primer 5′-TGATTTCAGTGCCTCTCCTG-3′. The founder was bred to C57BL/6NTac mice to propagate the strain.

## QUANTITATION AND STATISTICAL ANALYSIS

GraphPad Prism 9.1.1 were used for statistical analyses. Welch’s t-test, Mann-Whitney U test, or Wilson t-test was used as indicated in the figure and/ or legends. N values, replicate experiments are all located in figure legends. *P<0.05, **P<0.01, ***P <0.001.

## ACKNOWLEDGMENTS

This work was supported by the intramural Research Program of the NIH, National Eye Institute (project EY000184; R.R. Caspi). We gratefully acknowledge the editorial assistance of the NIH Fellows Editorial Board.Graphical abstract was created with BioRender.com.

## AUTHOR CONTRIBUTIONS

WJ and XX planned, performed and analyzed experiments and interpreted data. XX and WJ wrote the first manuscript draft. VN and JG analyzed ATAC-seq data. AG, ZP, AZ, JL, MJM, YJ, and RH performed experiments; YB and MC developed the C57BL/6-Trgv6em1Ybe (TCRγ6 KO) mice. RRC and XX conceptualized the study design and approaches and edited the manuscript. RRC conceived and supervised the study, reviewed and finalized the manuscript. All co-authors reviewed and approved the manuscript.

## REFERENCES

1. Li, X., Bechara, R., Zhao, J., McGeachy, M.J., and Gaffen, S.L. (2019). IL-17 receptor–based signaling and implications for disease. Nature Immunology 20, 1594–1602. 10.1038/s41590-019-0514-y.

2. Acosta-Rodriguez, E.V., Rivino, L., Geginat, J., Jarrossay, D., Gattorno, M., Lanzavecchia, A., Sallusto, F., and Napolitani, G. (2007). Surface phenotype and antigenic specificity of human interleukin 17–producing T helper memory cells. Nature Immunology 8, 639–646. 10.1038/ni1467.

3. Zhou, L., Ivanov, II, Spolski, R., Min, R., Shenderov, K., Egawa, T., Levy, D.E., Leonard, W.J., and Littman, D.R. (2007). IL-6 programs T(H)-17 cell differentiation by promoting sequential engagement of the IL-21 and IL-23 pathways. Nat Immunol 8, 967–974. 10.1038/ni1488.

4. Gomez-Rodriguez, J., Wohlfert, E.A., Handon, R., Meylan, F., Wu, J.Z., Anderson, S.M., Kirby, M.R., Belkaid, Y., and Schwartzberg, P.L. (2014). Itk-mediated integration of T cell receptor and cytokine signaling regulates the balance between Th17 and regulatory T cells. Journal of Experimental Medicine 211, 529–543. 10.1084/jem.20131459.

5. Sutton, C.E., Lalor, S.J., Sweeney, C.M., Brereton, C.F., Lavelle, E.C., and Mills, K.H. (2009). Interleukin-1 and IL-23 induce innate IL-17 production from gammadelta T cells, amplifying Th17 responses and autoimmunity. Immunity 31, 331–341. 10.1016/j.immuni.2009.08.001.

6. Duan, J., Chung, H., Troy, E., and Kasper, D.L. (2010). Microbial colonization drives expansion of IL-1 receptor 1-expressing and IL-17-producing gamma/delta T cells. Cell Host Microbe 7, 140–150. 10.1016/j.chom.2010.01.005.

7. St Leger, A.J., Hansen, A.M., Karauzum, H., Horai, R., Yu, C.R., Laurence, A., Mayer-Barber, K.D., Silver, P., Villasmil, R., Egwuagu, C., et al. (2018). STAT-3-independent production of IL-17 by mouse innate-like alphabeta T cells controls ocular infection. J Exp Med 215, 1079–1090. 10.1084/jem.20170369.

8. Sherlock, J.P., Joyce-Shaikh, B., Turner, S.P., Chao, C.C., Sathe, M., Grein, J., Gorman, D.M., Bowman, E.P., McClanahan, T.K., Yearley, J.H., et al. (2012). IL-23 induces spondyloarthropathy by acting on ROR-γt+ CD3+CD4-CD8-entheseal resident T cells. Nat Med 18, 1069–1076. 10.1038/nm.2817.

9. Edwards, S.C., Sutton, C.E., Ladell, K., Grant, E.J., McLaren, J.E., Roche, F., Dash, P., Apiwattanakul, N., Awad, W., Miners, K.L., et al. (2020). A population of proinflammatory T cells coexpresses αβ and γδ T cell receptors in mice and humans. J Exp Med 217. 10.1084/jem.20190834.

10. Knop, E., and Knop, N. (2005). The role of eye-associated lymphoid tissue in corneal immune protection. J Anat 206, 271–285. 10.1111/j.1469-7580.2005.00394.x.

11. Conti, H.R., Peterson, A.C., Brane, L., Huppler, A.R., Hernández-Santos, N., Whibley, N., Garg, A.V., Simpson-Abelson, M.R., Gibson, G.A., Mamo, A.J., et al. (2014). Oral-resident natural Th17 cells and γδ T cells control opportunistic Candida albicans infections. J Exp Med 211, 2075–2084. 10.1084/jem.20130877.

12. St Leger, A.J., Desai, J.V., Drummond, R.A., Kugadas, A., Almaghrabi, F., Silver, P., Raychaudhuri, K., Gadjeva, M., Iwakura, Y., Lionakis, M.S., and Caspi, R.R. (2017). An Ocular Commensal Protects against Corneal Infection by Driving an Interleukin-17 Response from Mucosal gammadelta T Cells. Immunity 47, 148–158 e145. 10.1016/j.immuni.2017.06.014.

13. de Paiva, C.S., St Leger, A.J., and Caspi, R.R. (2022). Mucosal immunology of the ocular surface. Mucosal Immunol 15, 1143–1157. 10.1038/s41385-022-00551-6.

14. Suryawanshi, A., Veiga-Parga, T., Rajasagi, N.K., Reddy, P.B., Sehrawat, S., Sharma, S., and Rouse, B.T. (2011). Role of IL-17 and Th17 cells in herpes simplex virus-induced corneal immunopathology. J Immunol 187, 1919–1930. 10.4049/jimmunol.1100736.

15. Gadjeva, M., Nagashima, J., Zaidi, T., Mitchell, R.A., and Pier, G.B. (2010). Inhibition of macrophage migration inhibitory factor ameliorates ocular Pseudomonas aeruginosa-induced keratitis. PLoS Pathog 6, e1000826. 10.1371/journal.ppat.1000826.

16. Akira, S., and Takeda, K. (2004). Toll-like receptor signalling. Nat Rev Immunol 4, 499–511. 10.1038/nri1391.

17. Reynolds, J.M., Pappu, B.P., Peng, J., Martinez, G.J., Zhang, Y., Chung, Y., Ma, L., Yang, X.O., Nurieva, R.I., Tian, Q., and Dong, C. (2010). Toll-like receptor 2 signaling in CD4(+) T lymphocytes promotes T helper 17 responses and regulates the pathogenesis of autoimmune disease. Immunity 32, 692–702. 10.1016/j.immuni.2010.04.010.

18. Martin, B., Hirota, K., Cua, D.J., Stockinger, B., and Veldhoen, M. (2009). Interleukin-17-producing gammadelta T cells selectively expand in response to pathogen products and environmental signals. Immunity 31, 321–330. 10.1016/j.immuni.2009.06.020.

19. Marks, K.E., Flaherty, S., Patterson, K.M., Stratton, M., Martinez, G.J., and Reynolds, J.M. (2021). Toll-like receptor 2 induces pathogenicity in Th17 cells and reveals a role for IPCEF in regulating Th17 cell migration. Cell Rep 35, 109303. 10.1016/j.celrep.2021.109303.

20. Cai, Y., Xue, F., Qin, H., Chen, X., Liu, N., Fleming, C., Hu, X., Zhang, H.G., Chen, F., Zheng, J., and Yan, J. (2019). Differential Roles of the mTOR-STAT3 Signaling in Dermal gammadelta T Cell Effector Function in Skin Inflammation. Cell Rep 27, 3034–3048 e3035. 10.1016/j.celrep.2019.05.019.

21. Lopes, N., McIntyre, C., Martin, S., Raverdeau, M., Sumaria, N., Kohlgruber, A.C., Fiala, G.J., Agudelo, L.Z., Dyck, L., Kane, H., et al. (2021). Distinct metabolic programs established in the thymus control effector functions of gammadelta T cell subsets in tumor microenvironments. Nat Immunol 22, 179–192. 10.1038/s41590-020-00848-3.

22. Oliveira-Nascimento, L., Massari, P., and Wetzler, L.M. (2012). The Role of TLR2 in Infection and Immunity. Front Immunol 3, 79. 10.3389/fimmu.2012.00079.

23. Mokuno, Y., Matsuguchi, T., Takano, M., Nishimura, H., Washizu, J., Ogawa, T., Takeuchi, O., Akira, S., Nimura, Y., and Yoshikai, Y. (2000). Expression of toll-like receptor 2 on gamma delta T cells bearing invariant V gamma 6/V delta 1 induced by Escherichia coli infection in mice. J Immunol 165, 931–940. 10.4049/jimmunol.165.2.931.

24. Heilig, J.S., and Tonegawa, S. (1986). Diversity of murine gamma genes and expression in fetal and adult T lymphocytes. Nature 322, 836–840. 10.1038/322836a0.

25. O’Brien, R.L., and Born, W.K. (2020). Two functionally distinct subsets of IL-17 producing γd T cells. Immunol Rev 298, 10–24. 10.1111/imr.12905.

26. Shibata, K., Yamada, H., Hara, H., Kishihara, K., and Yoshikai, Y. (2007). Resident Vdelta1+ gammadelta T cells control early infiltration of neutrophils after Escherichia coli infection via IL-17 production. J Immunol 178, 4466–4472. 10.4049/jimmunol.178.7.4466.

27. Wei, Y.L., Han, A., Glanville, J., Fang, F., Zuniga, L.A., Lee, J.S., Cua, D.J., and Chien, Y.H. (2015). A Highly Focused Antigen Receptor Repertoire Characterizes γδ T Cells That are Poised to Make IL-17 Rapidly in Naive Animals. Front Immunol 6, 118. 10.3389/fimmu.2015.00118.

28. Roark, C.L., Aydintug, M.K., Lewis, J., Yin, X., Lahn, M., Hahn, Y.S., Born, W.K., Tigelaar, R.E., and O’Brien, R.L. (2004). Subset-specific, uniform activation among V gamma 6/V delta 1+ gamma delta T cells elicited by inflammation. J Leukoc Biol 75, 68–75. 10.1189/jlb.0703326.

29. Wang, X., Zhang, Y., Yang, X.O., Nurieva, R.I., Chang, S.H., Ojeda, S.S., Kang, H.S., Schluns, K.S., Gui, J., Jetten, A.M., and Dong, C. (2012). Transcription of Il17 and Il17f is controlled by conserved noncoding sequence 2. Immunity 36, 23–31. 10.1016/j.immuni.2011.10.019.

30. Akimzhanov, A.M., Yang, X.O., and Dong, C. (2007). Chromatin remodeling of interleukin-17 (IL-17)-IL-17F cytokine gene locus during inflammatory helper T cell differentiation. J Biol Chem 282, 5969–5972. 10.1074/jbc.C600322200.

31. Okamoto, K., Iwai, Y., Oh-Hora, M., Yamamoto, M., Morio, T., Aoki, K., Ohya, K., Jetten, A.M., Akira, S., Muta, T., and Takayanagi, H. (2010). IkappaBzeta regulates T(H)17 development by cooperating with ROR nuclear receptors. Nature 464, 1381–1385. 10.1038/nature08922.

32. Hayes, S.M., Li, L., and Love, P.E. (2005). TCR Signal Strength Influences αβ/γδ Lineage Fate. Immunity 22, 583–593. 10.1016/j.immuni.2005.03.014.

33. Yoshida, H., Lareau, C.A., Ramirez, R.N., Rose, S.A., Maier, B., Wroblewska, A., Desland, F., Chudnovskiy, A., Mortha, A., Dominguez, C., et al. (2019). The cis-Regulatory Atlas of the Mouse Immune System. Cell 176, 897-912.e820. 10.1016/j.cell.2018.12.036.

34. Raud, B., Roy, D.G., Divakaruni, A.S., Tarasenko, T.N., Franke, R., Ma, E.H., Samborska, B., Hsieh, W.Y., Wong, A.H., Stüve, P., et al. (2018). Etomoxir Actions on Regulatory and Memory T Cells Are Independent of Cpt1a-Mediated Fatty Acid Oxidation. Cell Metabolism 28, 504-515.e507. 10.1016/j.cmet.2018.06.002.

35. Majumder, S., Amatya, N., Revu, S., Jawale, C.V., Wu, D., Rittenhouse, N., Menk, A., Kupul, S., Du, F., Raphael, I., et al. (2019). IL-17 metabolically reprograms activated fibroblastic reticular cells for proliferation and survival. Nat Immunol 20, 534–545. 10.1038/s41590-019-0367-4.

36. Wilharm, A., Tabib, Y., Nassar, M., Reinhardt, A., Mizraji, G., Sandrock, I., Heyman, O., Barros-Martins, J., Aizenbud, Y., Khalaileh, A., et al. (2019). Mutual interplay between IL-17-producing gammadeltaT cells and microbiota orchestrates oral mucosal homeostasis. Proceedings of the National Academy of Sciences of the United States of America 116, 2652–2661. 10.1073/pnas.1818812116.

37. Lu, Y., Cao, X., Zhang, X., and Kovalovsky, D. (2015). PLZF Controls the Development of Fetal-Derived IL-17+Vgamma6+ gammadelta T Cells. J Immunol 195, 4273–4281. 10.4049/jimmunol.1500939.

38. Marchitto, M.C., Dillen, C.A., Liu, H., Miller, R.J., Archer, N.K., Ortines, R.V., Alphonse, M.P., Marusina, A.I., Merleev, A.A., Wang, Y., et al. (2019). Clonal Vγ6(+)Vd4(+) T cells promote IL-17-mediated immunity against Staphylococcus aureus skin infection. Proceedings of the National Academy of Sciences of the United States of America 116, 10917–10926. 10.1073/pnas.1818256116.

39. Ciofani, M., Madar, A., Galan, C., Sellars, M., Mace, K., Pauli, F., Agarwal, A., Huang, W., Parkhurst, C.N., Muratet, M., et al. (2012). A validated regulatory network for Th17 cell specification. Cell 151, 289–303. 10.1016/j.cell.2012.09.016.

40. Omenetti, S., Bussi, C., Metidji, A., Iseppon, A., Lee, S., Tolaini, M., Li, Y., Kelly, G., Chakravarty, P., Shoaie, S., et al. (2019). The Intestine Harbors Functionally Distinct Homeostatic Tissue-Resident and Inflammatory Th17 Cells. Immunity 51, 77-89.e76. 10.1016/j.immuni.2019.05.004.

41. Di Luccia, B., Gilfillan, S., Cella, M., Colonna, M., and Huang, S.C. (2019). ILC3s integrate glycolysis and mitochondrial production of reactive oxygen species to fulfill activation demands. J Exp Med 216, 2231–2241. 10.1084/jem.20180549.

42. Ivanov, II, Atarashi, K., Manel, N., Brodie, E.L., Shima, T., Karaoz, U., Wei, D., Goldfarb, K.C., Santee, C.A., Lynch, S.V., et al. (2009). Induction of intestinal Th17 cells by segmented filamentous bacteria. Cell 139, 485–498. 10.1016/j.cell.2009.09.033.

43. Honda, K., and Littman, D.R. (2016). The microbiota in adaptive immune homeostasis and disease. Nature 535, 75–84. 10.1038/nature18848.

44. Corces, M.R., Trevino, A.E., Hamilton, E.G., Greenside, P.G., Sinnott-Armstrong, N.A., Vesuna, S., Satpathy, A.T., Rubin, A.J., Montine, K.S., Wu, B., et al. (2017). An improved ATAC-seq protocol reduces background and enables interrogation of frozen tissues. Nat Methods 14, 959–962. 10.1038/nmeth.4396.

45. Chen, S., Zhou, Y., Chen, Y., and Gu, J. (2018). fastp: an ultra-fast all-in-one FASTQ preprocessor. Bioinformatics 34, i884–i890. 10.1093/bioinformatics/bty560.

46. Langmead, B., and Salzberg, S.L. (2012). Fast gapped-read alignment with Bowtie 2. Nat Methods 9, 357–359. 10.1038/nmeth.1923.

47. Gaspar, J.M. (2021). Genrich: detecting sites of genomic enrichment. https://github.com/jsh58/Genrich.

48. Yu, G., Wang, L.G., and He, Q.Y. (2015). ChIPseeker: an R/Bioconductor package for ChIP peak annotation, comparison and visualization. Bioinformatics 31, 2382–2383. 10.1093/bioinformatics/btv145.

49. Love, M.I., Huber, W., and Anders, S. (2014). Moderated estimation of fold change and dispersion for RNA-seq data with DESeq2. Genome Biol 15, 550. 10.1186/s13059-014-0550-8.

50. Karolchik, D., Baertsch, R., Diekhans, M., Furey, T.S., Hinrichs, A., Lu, Y.T., Roskin, K.M., Schwartz, M., Sugnet, C.W., Thomas, D.J., et al. (2003). The UCSC Genome Browser Database. Nucleic Acids Res 31, 51–54. 10.1093/nar/gkg129.

