## Supplementary material for "TLR2 Supports γδ T cell IL-17A Response to ocular surface commensals by Metabolic Reprogramming": All supplementary Figures

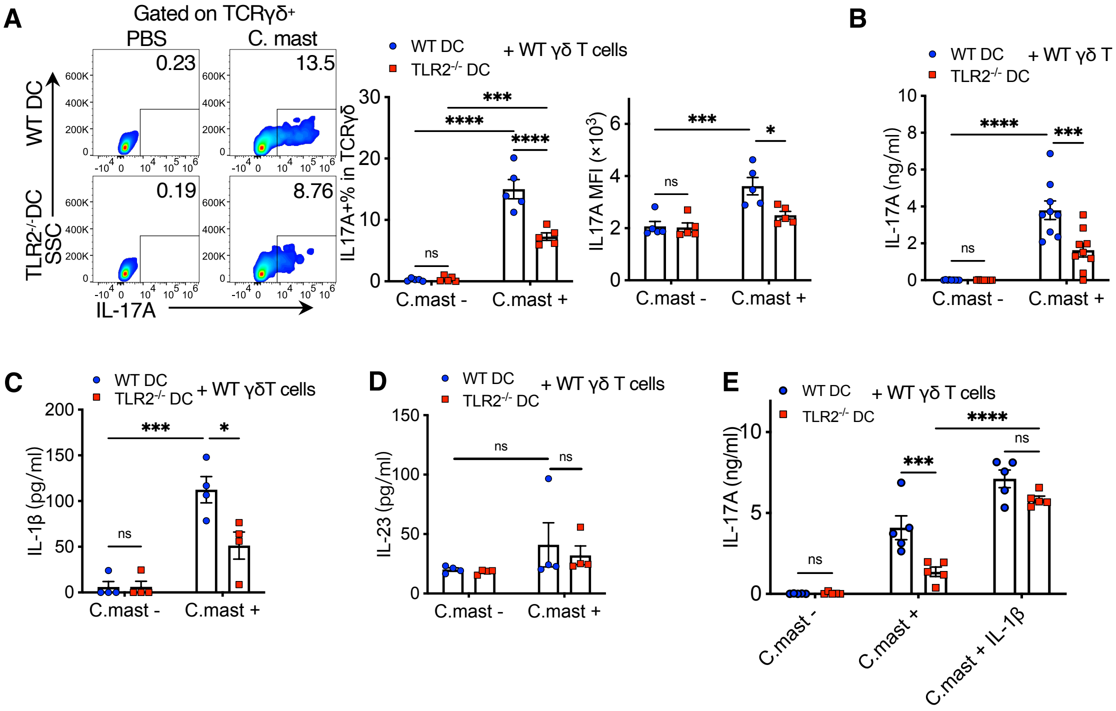


**Figure S1 (refers to Figure 2). TLR2 expressed by DC drives IL-1β needed to support the γδ T cell IL-17A response to *C. mast***

**(A-E)** WT γδ T cells were isolated from WT mice and co-cocultured with WT or TLR2$^{-/-}$ CD11c+ dendritic cells for 72 hours (γδ T cells: $2\times10^4$; CD11c+ cells: $1\times10^5$ ) with/ without heat-killed *C. mast*. Supernatants and cells were collected for ELISA and flow cytometry experiments.
**(A)** Representative FACS plots and bar graphs showing the percentage and mean fluorescence index (MFI) of IL-17A in γδ T cells from WT or TLR2$^{-/-}$ DC co-culture. N=5. Combined data from 3 experiments.
**(B-D)** Bar graphs presenting the levels of IL-17A **(B)**, IL-1β **(C)**, and IL-23 **(D)** proteins in the supernatants from WT or TLR2$^{-/-}$ DC co-culture system. Data was combined from 4 experiments (N=9) for (**B**) and 2 experiments (N=4) for **(C-D)**.
**(E)** IL-1β (100ng/ml) was added to the co-culture system for 72h, and the IL-17A level was measured by ELISA. N=5. Combined data from 3 experiments.
Statistical significance was determined by Two way ANOVA (A-E). (A-E) Bars represent mean ± SEM with *P<0.05, **P<0.01, ***P <0.001, **** P < 0.0001.

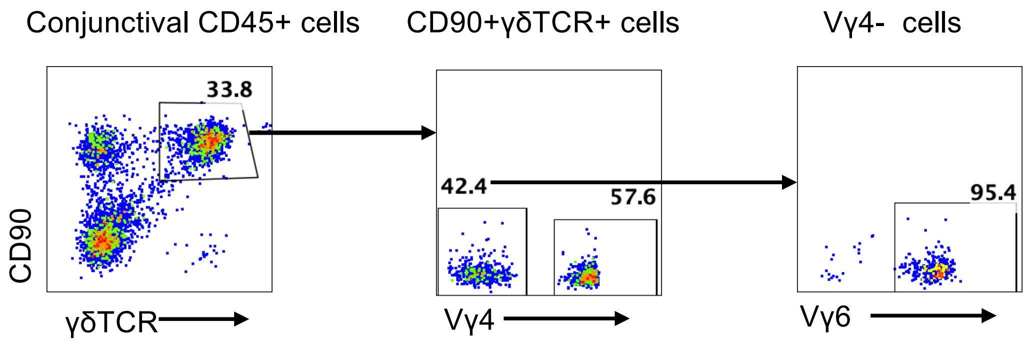


**Figure S2 (refers to Figure 4).** Conjunctival Vγ4 – cells  expressed Vγ6 TCR.

Conjunctival cells from WT mice were stained with anti-Vγ4 and anti-Vγ6 antibodies and analyzed by flow cytometry.

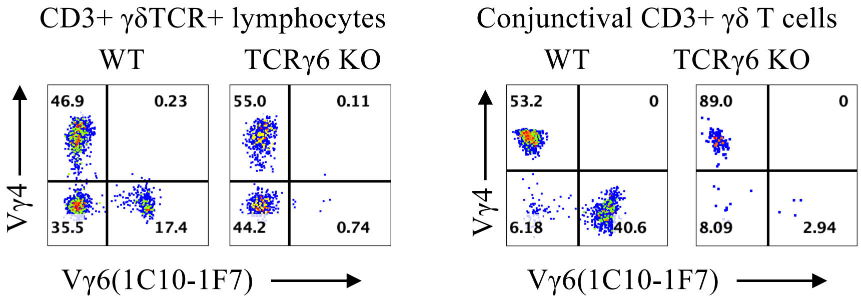


**Figure S3 (refers to Figure 4).** TCRγ6 KO mice are confirmed by the absence of Vγ6 cells in the lymph node and conjunctiva.

Lymphocytes from eye draining LNs and conjunctival cells from WT and TCRγ6 KO mice were stained with the indicated antibodies and analyzed by flow cytometry.

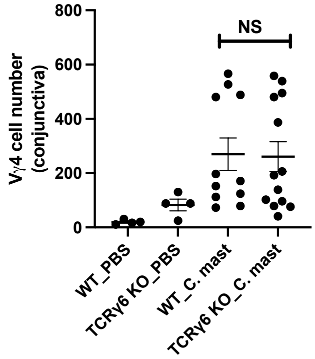


**Figure S4 (refers to Figure 4).** The number of Vγ4 cells in conjunctivae was comparable between WT and TCRγ6 KO mice.

WT and TCRγ6 KO mice were inoculated every other day with either *C. mast* ($5 \times 10^5$ CFU) or PBS for three inoculations. Seven days after inoculation, conjunctivae were analyzed for the number of Vγ4 cells. Each dot represent one animal. Significance was determined by two-way ANOVA.

**A** Gating strategy of Vγ6 cells

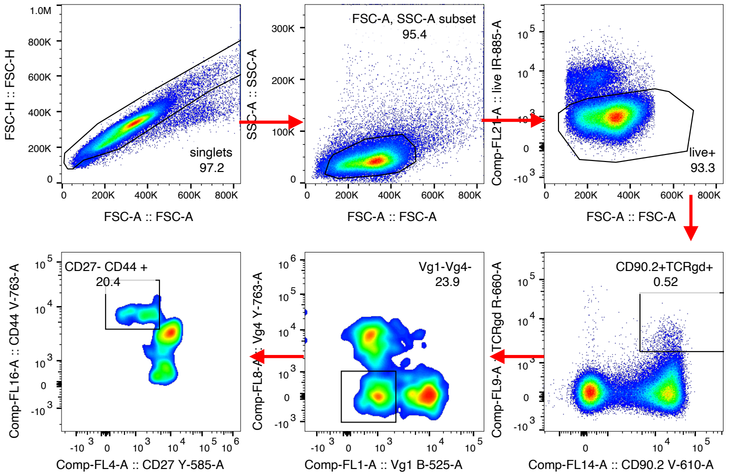


**B** ATAC-seq (ImmGen)

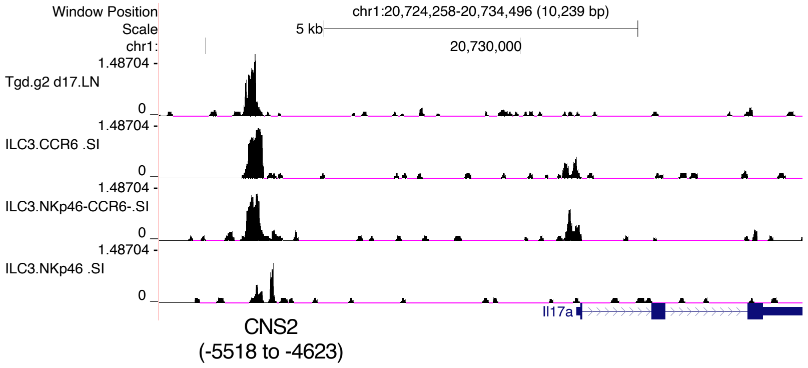


**C** T7 endonuclease assay

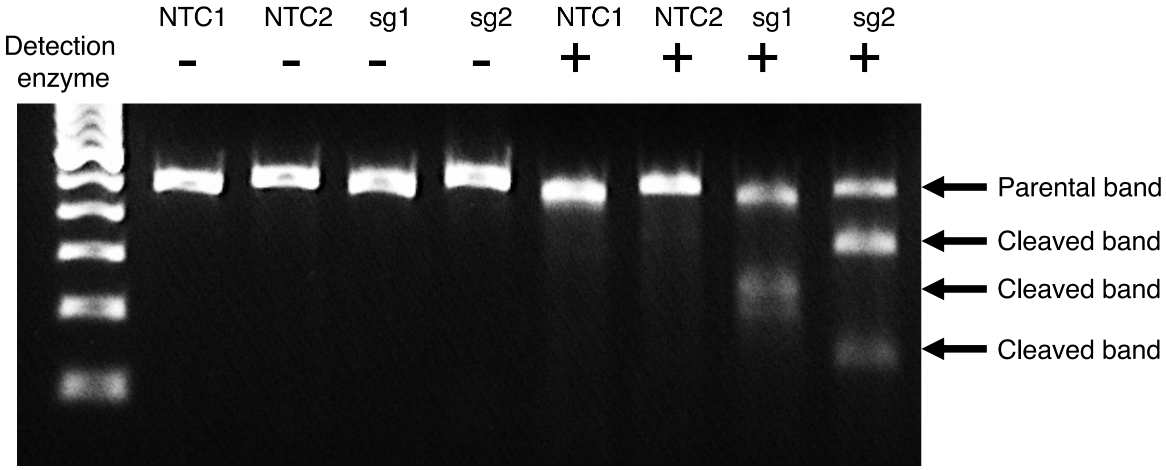


**Figure S5 (refers to Figure 5). Decreased IκBζ is causally related to deficient TLR2-dependent IL-17A production in Vγ6 cells**

**(A)** Representative FACS plots showing CD27$^{-}$CD44$^{high}$ Vγ1-Vγ4- gating strategy from eye draining lymph nodes of *C. mast*$^{-}$ or *C. mast*$^{+}$ TLR2$^{-/-}$ mice for RNA-seq and ATAC-seq.
**(B)** The UCSC genome browser-depicted traces, to the same scale, for the Il17a genomic region of the indicated cell subsets using ATAC-seq under the ImmGen Program.
**(C)** Vγ6 cells were transfected with CRISPR ribonucleoprotein complexes targeting different regions of Nfkbiz (sg1 and sg2) or a non-targeting control(NTC) that does not target the mouse reference genomes. PCR products specific to the Nfkbiz target region were analyzed by the T7 E1 mismatch cleavage assay. Cleaved bands indicate successful CRISPR editing.

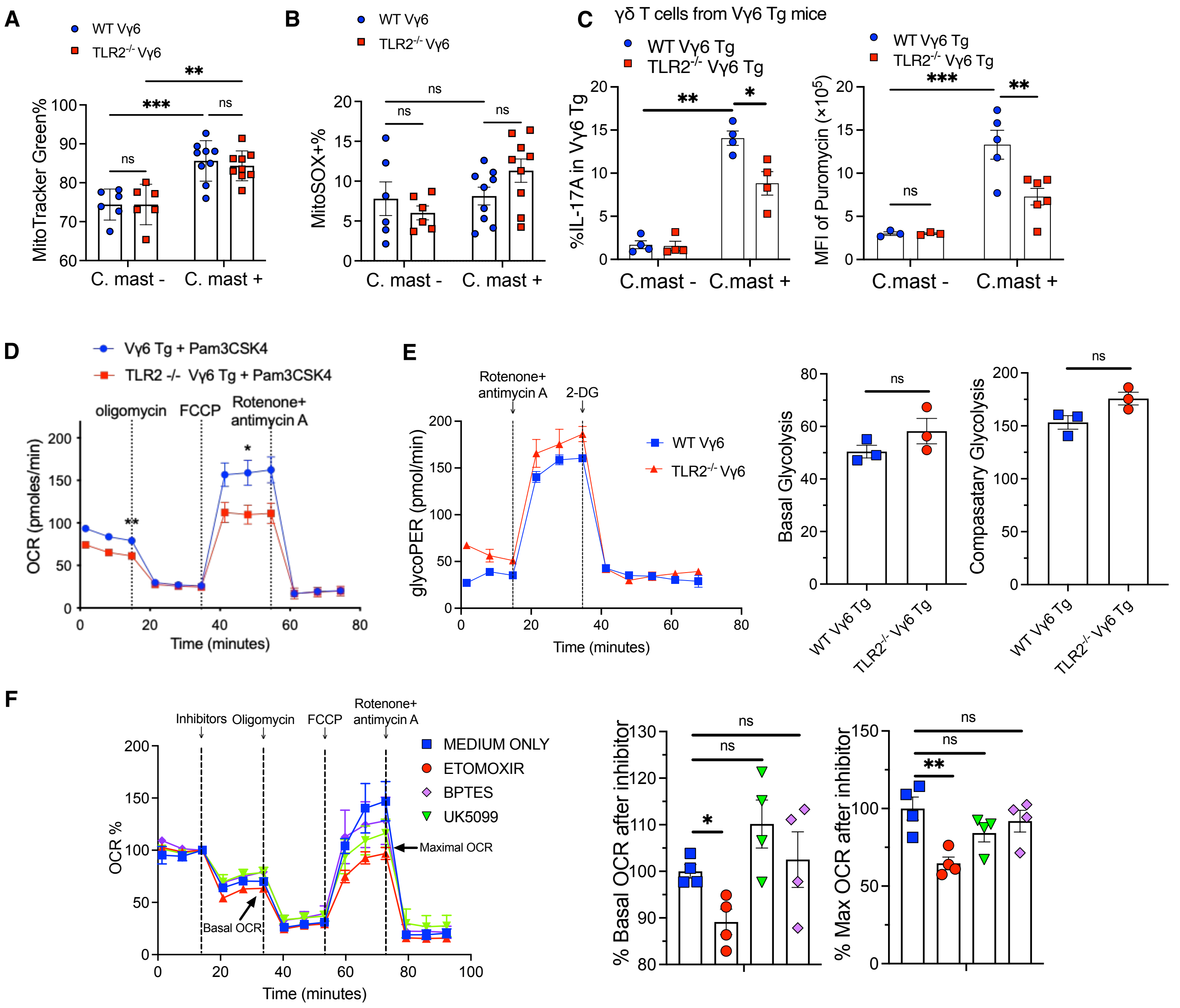


**Figure S6 (refers to Figure 6). TLR2 deficiency Vγ6 T cells do not impair glycolysis**

γδ T cells were collected from secondary lymphoid tissues of TLR2 -/- or WT mice and subjected to Seahorse Metabolic profiling.

**(A-B)** FACS plots and bar graphs showing the percentage of MitoTrack Green **(A)** and MitoSOX (mitochondrial Superoxide Indicator **(B)** in Vγ6 cells from WT and TLR2$^{-/-}$ mice. N= 3–6. Combined data from 2 independent experiments.

**(C)** Vγ6 cells isolated from WT and TLR2$^{-/-}$ Vγ6 Tg mice were co-cocultured with WT CD11c$^{+}$ dendritic cells for 72 hours (γδ T cells: $2\times10^4$; CD11c+ cells: $1\times10^5$ ) with or without heat-killed *C. mast*. The percentage of IL-17A$^{+}$ cells (left plot) was measured by flow cytometry after staining for anti-IL17A. For ATP measurement, puromycin was added to the culture for the last 1 hour and then Vγ6 cells were stained with anti-puromycin. The MFI of incorporated puromycin reflects ATP production. N= 3–6.

**(D)** Vγ6 cells were sorted from WT and TLR2$^{-/-}$ Vγ6 Tg mice treated with Pam3CSK4 for 3 days and then were subjected to the Seahorse Mito stress test. Representative data from one of 3 independent experiments.

**(E)** Vγ6 T cells isolated from WT and TLR2$^{-/}$ *C. mast*+ Vγ6 transgenic mice were subjected to GlycoPER assay. Bar graphs show basal Glycolysis and compensatory glycolysis of WT and TLR2$^{-/-}$ Vγ6 cells. N= 3. Representative data from one of 2 independent experiments.

**(F)** Vγ6 cells in *C. mast*$^{+}$ Vγ6 transgenic mice were sorted and performed seahorse substrate oxidation stress test. Inhibitors (ETOMOXIR, UK5099, and BPTES) were injected before oligomycin treatment. Percentage of basal OCR after inhibitors injection and the percentage of maximal OCR after inhibitor injection were calculated and displayed in bar plots. N= 4. Representative data from one of 2 independent experiments.
Significance was determined by Two way ANOVA in (A-C) or Welch's t-test (E-F).

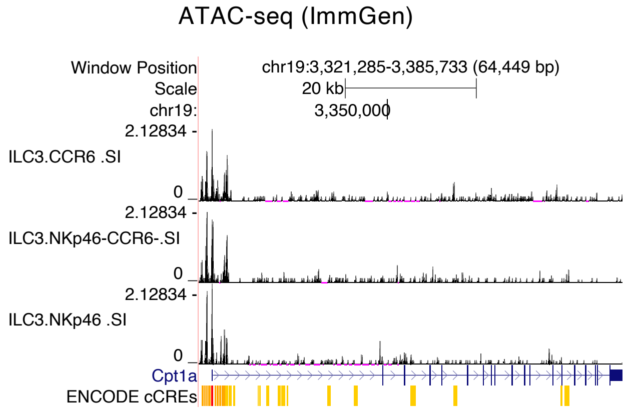


**Figure S7 (refers to Figure 7)**

UCSC genome browser output depicting ATAC-seq traces for the promoter and enhancer regions (ENCODE cCREs) of *Cpt1a* in indicated IL-17A producing immune cells sequenced by ImmGen program.

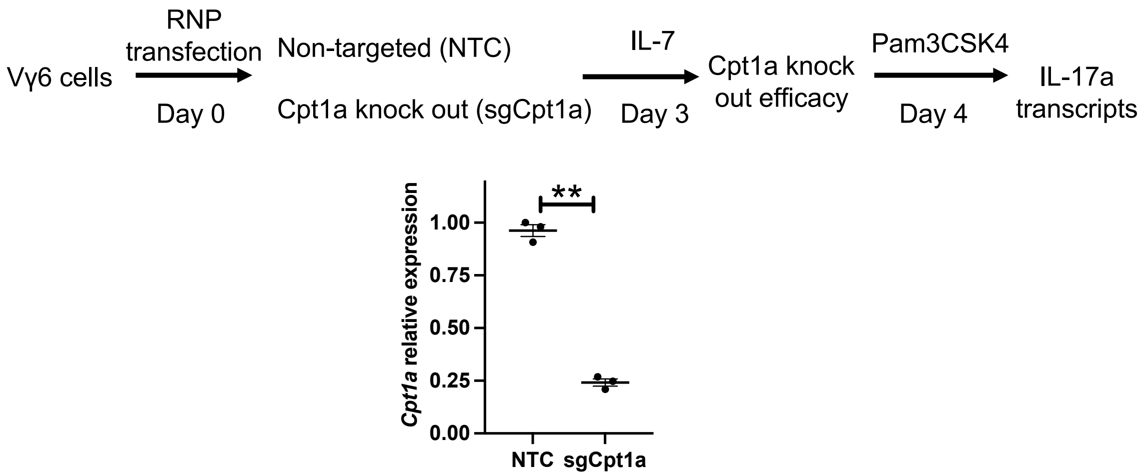


**Figure S8 (refers to Figure 7) Generation of Cpt1a-deficient Vγ6 cells using CRISPR**

Vγ6 cells were transfected with CRISPR ribonucleoprotein complexes targeting different Cpt1a loci. 3 days post-transfection, RNA was extracted from CRISPR-edited samples and analyzed for *Cpt1a* transcripts by real-time PCR. Each dot represents one experiment. Significance was determined by Welch’s t-test.
